# Molecular determinants of antibody-mediated priming to enhance detection of ctDNA

**DOI:** 10.64898/2026.01.27.701975

**Authors:** Shervin Tabrizi, Claire Sullivan, Kasturi Chakraborty, Carmen Martin-Alonso, Zhenyi An, Lily Gao, Daniel M. Kim, Savan K. Patel, Timothy Blewett, Justin Rhoades, Ruolin Liu, Sahil Patel, Kan Xiong, Andjela Crnjac, Sangeeta N. Bhatia, Viktor A. Adalsteinsson, J. Christopher Love

## Abstract

Liquid biopsies can enable cancer detection and monitoring yet remain limited by low concentrations of ctDNA. To address this limitation, we previously introduced a monoclonal antibody (mAb) “priming agent” that transiently increased the concentration of ctDNA in blood. Here, we investigated the molecular features that drive this effect. In a panel of novel mAbs that bound cfDNA, both those targeting dsDNA and those targeting mononucleosomes increased the concentration of ctDNA. mAbs with high avidity to dsDNA performed best, suggesting dsDNA as the key binding target. One agent preserved short, biologically informative molecules of cfDNA typically depleted at baseline. The Fc domain was dispensable, as F(ab’)2 fragments retained priming activity. Leveraging these insights, we engineered single-chain agents using the dsDNA binding domain sso7d, expanding priming strategies beyond immunoglobulins. This study identifies the molecular features and general design principles for mAb-based priming agents to enhance recovery and detection of ctDNA.

**Significance:** Low concentration of ctDNA limits the sensitivity of liquid biopsies in many applications. This study shows how engineered antibodies and other dsDNA-binding molecules can inhibit clearance of ctDNA from the bloodstream and increase concentration of ctDNA in a blood draw, and identifies the key features that enable this activity. Modulation of cfDNA in the bloodstream using these agents can increase recovery of ctDNA and improve the performance of liquid biopsies for cancer detection.

## Introduction

Detection of circulating tumor DNA (ctDNA) via liquid biopsies can enable early detection of cancer, monitoring for minimal residual disease (MRD), and identification of clinically actionable mutations in plasma to inform therapy selection.^1,2^ Low quantities of ctDNA in plasma in settings of early-stage cancer and MRD have prompted the development of ultrasensitive assays that comprehensively profile molecules of cfDNA to detect rare fragments of ctDNA.^3–5^ Despite increasingly deeper and broader profiling of cfDNA, current state-of-the-art tests are limited when disease burden is low.^6^ Due to exceedingly low amounts of ctDNA in these settings, molecules of ctDNA are often physically absent in blood draws, making their detection impossible.^1,7^ Increasing total quantities of cfDNA used in these assays can overcome this limitation and improve sensitivity.^6,8,9^

We previously reported on injectable priming agents that can improve the recovery of ctDNA by inhibiting clearance of cfDNA from the bloodstream.^9^ An immunologically-silenced monoclonal antibody (mAb) against dsDNA (“aST3”) inhibited clearance of cfDNA, improved recovery of ctDNA, enhanced tumor mutational profiling from plasma, and improved the sensitivity of ctDNA-based tests.^9^ These results established proof-of-concept that a monoclonal antibody can improve ctDNA recovery. Yet, it remained unclear whether the priming effect would generalize to other mAbs, and whether the epitope, avidity, or the Fc domain play roles in mediating the priming effect. Answering these questions is critical for understanding the generalizability of cfDNA binders as a priming strategy and for informing strategies to screen and develop future priming agents.

To better understand mAbs as priming agents, we generated and evaluated novel engineered mAbs and their derivatives using a combination of *in vitro* biophysical and biochemical assays as well as *in vivo* models of cfDNA and ctDNA. We found multiple mAbs beyond aST3, including those with murine- and human-derived variable regions, that showed priming activity in tumor-bearing mice. Some mAbs specifically bound intact mononunuclosomes (MNs) whereas others bound dsDNA, both free and in MN. Specificity and avidity were both associated with the priming effect. Some priming agents preferentially protected short, biologically-informative fragments of cfDNA in circulation. The Fc-domain was dispensable for priming, suggesting that agents with more rapid clearance from plasma could still elicit a priming effect. We used our insights into the molecular determinants of mAb-mediated priming to design new molecules and demonstrated their ability to act as priming agents, supporting the generalizability of these principles.

## Results

### Engineered mAbs as priming agents recognize different targets and exhibit a range of binding kinetics

We designed and expressed a panel of 14 mAbs on the same backbone as aST3 comprising a murine IgG2a Fc with L234A/L235A/P329G mutations to silence effector function (**Fig. 1A, Table S1**).^10^ These mAbs were expressed using reported variable region (VH and VL) sequences of mAbs against components of cfDNA, including dsDNA (12 mAbs) and histones/MNs (2 mAbs).^11–15^ The mAbs used a variety of germline genes and showed significant sequence diversity (**Table S2, Fig. S1**). Data from electrophoretic mobility shift assays (EMSA) showed that four of the mAbs bound dsDNA in two conformations, as free dsDNA and as dsDNA in MNs. These four mAbs are hereafter referred to as DNA1, DNA2, DNA3, DNA4. In contrast, four mAbs bound intact MN only and showed no detectable binding to free dsDNA (hereafter referred to as MN1, MN2, MN3, MN4) (**Fig. 1B**). Intriguingly, all four mAbs that only bound MN had previously been reported to bind dsDNA on ELISA assays prepared using calf thymus DNA,^11^ suggesting potential chromatin contamination in calf thymus-derived DNA preparation in previous assays. Four of the remaining mAbs were reported in the literature to bind dsDNA and two were reported to bind histones/MNs; these six showed no detectable binding to dsDNA or MN.

**Figure 1:**
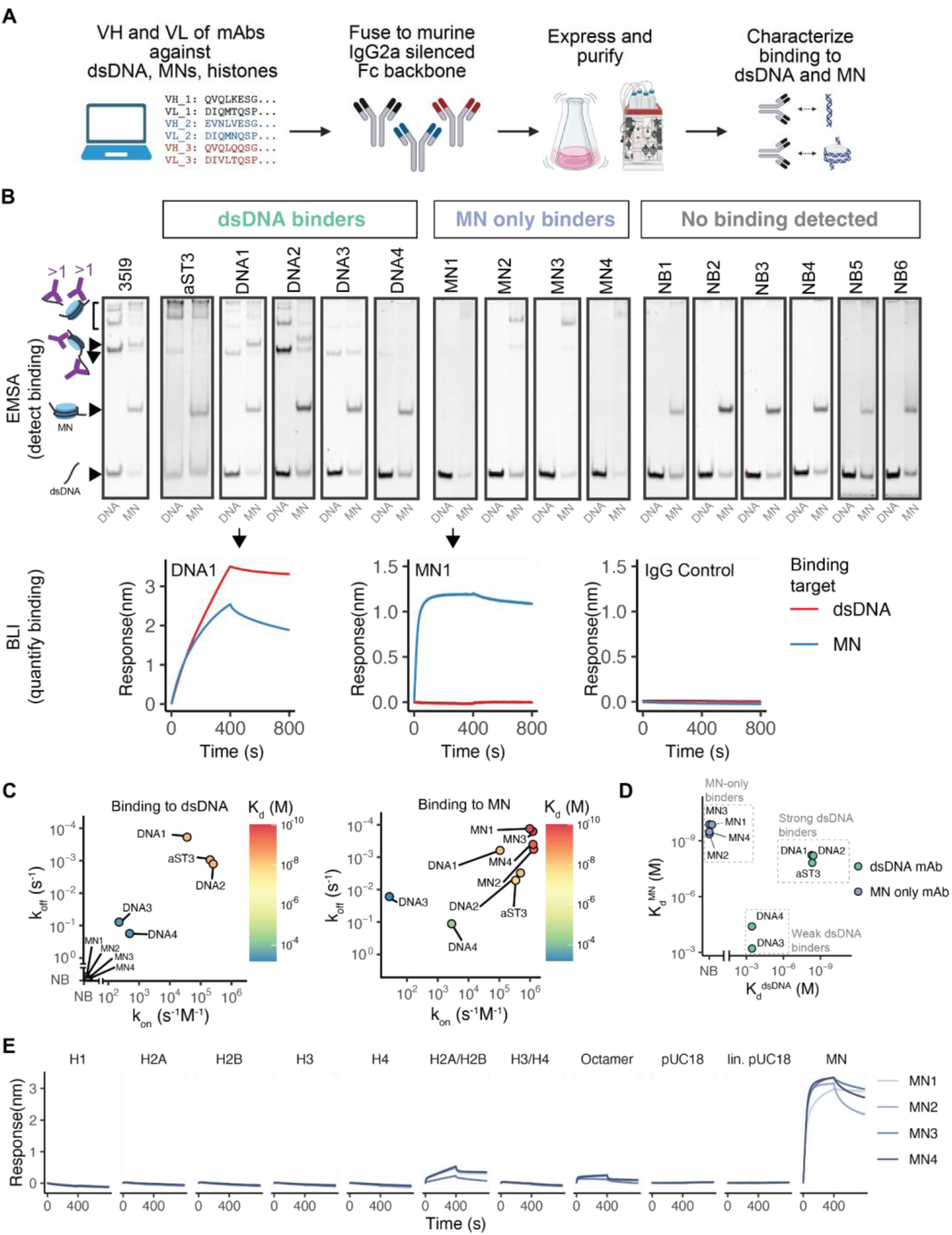
mAbs against cfDNA recognize dsDNA or intact mononucleosomes and exhibit a range of binding kinetics. A) Schematic of approach to design, express, and characterize a panel of immunologically-silenced mAbs against cfDNA components (dsDNA, mononucleosomes, histones). Priming agent mAbs were generated using VH and VL amino acid sequences of mAbs that bind cfDNA components fused to a murine IgG2a backbone with L234A/L235A/P329G silencing mutations. B) EMSA of mAbs binding to free dsDNA or MNs, and quantification of binding interaction with biolayer interferometry. Free dsDNA binding was determined by the presence of shifted bands in the DNA lane. MN-only binding was determined by shift or disappearance of the MN band in the MN lane and no evidence of a shifted band in the DNA lane. For mAbs that bound free dsDNA or MN, the binding interaction was subsequently quantified with BLI. C) Measurement of binding kinetics (k_on_ and k_off_) and avidity (apparent K_d_) to free dsDNA (left) and MN (right) of intact mAbs. D) Comparison of K_d_ of dsDNA binders and MN binders to free dsDNA and MN. E) Interaction of MN binders with individual components of mononucleosomes and different dsDNA topologies, including individual histones, H2A/H2B dimer, H3/H4 tetramer, H2A/H2B/H3/H4 octamer, bent dsDNA (supercoiled pUC18), linear dsDNA (linearized pUC18, lin. pUC18), and intact MN (histone octamer + 147bp dsDNA). NB - no binding.

For those mAbs that showed any binding activity on EMSA, we then measured their binding kinetics to free dsDNA and MN using biolayer interferometry (BLI) (**Fig. 1B, Table S3**). Monogamous bivalency – binding of both variable domains to the same molecule of dsDNA – is important for the binding interaction between antibodies and dsDNA and single F(ab) domains typically have little to no detectable binding to dsDNA.^16,17^ We therefore estimated the apparent K_d_ for full-length, bivalent IgG mAbs to include effects of avidity from monogamous bivalency. Avidities to dsDNA (K_d_^dsDNA^) varied significantly among the mAbs binding dsDNA, with two showing comparable K_d_ ^dsDNA^ to aST3 (aST3: 4.6nM, DNA1: 5.2nM, DNA2: 4.9nM), but with a wide range of association (k_on_) and dissociation (k_off_) rates. Two showed significantly worse avidity (DNA3: 0.35 mM, DNA4: 0.36 mM) (**Fig. 1C,D**). Avidity of the dsDNA binders to MNs was slightly weaker than to free dsDNA. In contrast, the mAbs that bound MN only showed no detectable binding to free dsDNA, but very high avidity to MN (all with K_d_^MN^ < 1nM), exceeding the avidity of dsDNA binding mAbs to MN (**Fig. 1C,D**). Comparing K_d_^dsDNA^ and K_d_^MN^ highlighted three groups of agents in our panel: 1) “Strong dsDNA binders,” which bound free dsDNA and dsDNA in MN with high avidity (aST3, DNA1, DNA2; K_d_^dsDNA^ and K_d_^MN^ <100 nM), 2), “Weak dsDNA binders,” which bound free dsDNA and dsDNA in MN with low avidity (K_d_^dsDNA^ and K_d_^MN^ > 0.1mM) and 3), and “MN-only binders,” which showed no binding activity to free dsDNA but very high avidity to MN (K_d_^MN^ < 1nM) (**Fig. 1D**). Given that the MN-only binders showed no binding to free dsDNA but high avidity to MN, we further investigated whether they bound individual substructures within the MN, or if they only recognized an intact MN comprising dsDNA wrapped around a histone octamer. The MN-only binders showed no binding to histones H1, H2A, H2B, H3, H4 and H3/H4 tetramer and very weak binding to H2A/H2B dimers and H2A/H2B/H3/H4 octamer (**Fig. 1E**). Furthermore, they showed no detectable binding to bent dsDNA in supercoiled pUC18 plasmid or to linearized pUC18. They maintained high avidity binding to intact MN, consisting of dsDNA and the histone octamer (**Fig. 1E**). These findings suggest that the MN-only binders specifically bind intact MN with high avidity.

### Impact of mAb priming agents on clearance of dsDNA and MN

We next characterized the interaction of our mAbs with cfDNA and their impact on clearance of cfDNA. Cell-free DNA is degraded by nucleases and cleared by organs such as the liver and kidney.^18–20^ To evaluate the ability of mAbs as priming agents to protect DNA from degradation by nucleases, we measured the degradation of DNA in the presence of DNase at an activity level of 0.2U/mL (comparable to plasma^21–23)^ with or without mAbs over time. Antibodies that bound dsDNA inhibited the degradation of free dsDNA over time, whereas antibodies that only bound MN had no effect on the degradation of free dsDNA, suggesting that physical interaction with free dsDNA is necessary to protect it from degradation by nucleases (**Fig. 2A, left**). In contrast to free dsDNA in the absence of mAbs, dsDNA wrapped around histones in MNs (“MN-bound dsDNA”) was significantly more resistant to degradation by nucleases, reflecting the protective effect of histones on degradation by nucleases for MN-bound dsDNA (**Fig. 2A, right**). Antibodies that bound dsDNA still inhibited degradation of MN-bound dsDNA, whereas antibodies that bound MNs did not provide any apparent protection, despite the MN-only mAbs all having higher avidity to MN compared to the dsDNA mAbs (**Fig. 2A**, **right**). Therefore, degradation by nucleases results in rapid degradation of free dsDNA but not MN-bound dsDNA in MN, likely due to the protective effect of histones. Furthermore, mAbs that directly bind dsDNA were superior in protecting both free dsDNA and MN-bound dsDNA from degradation, suggesting that direct interaction between mAb and dsDNA is important for the protective effect.

**Figure 2:**
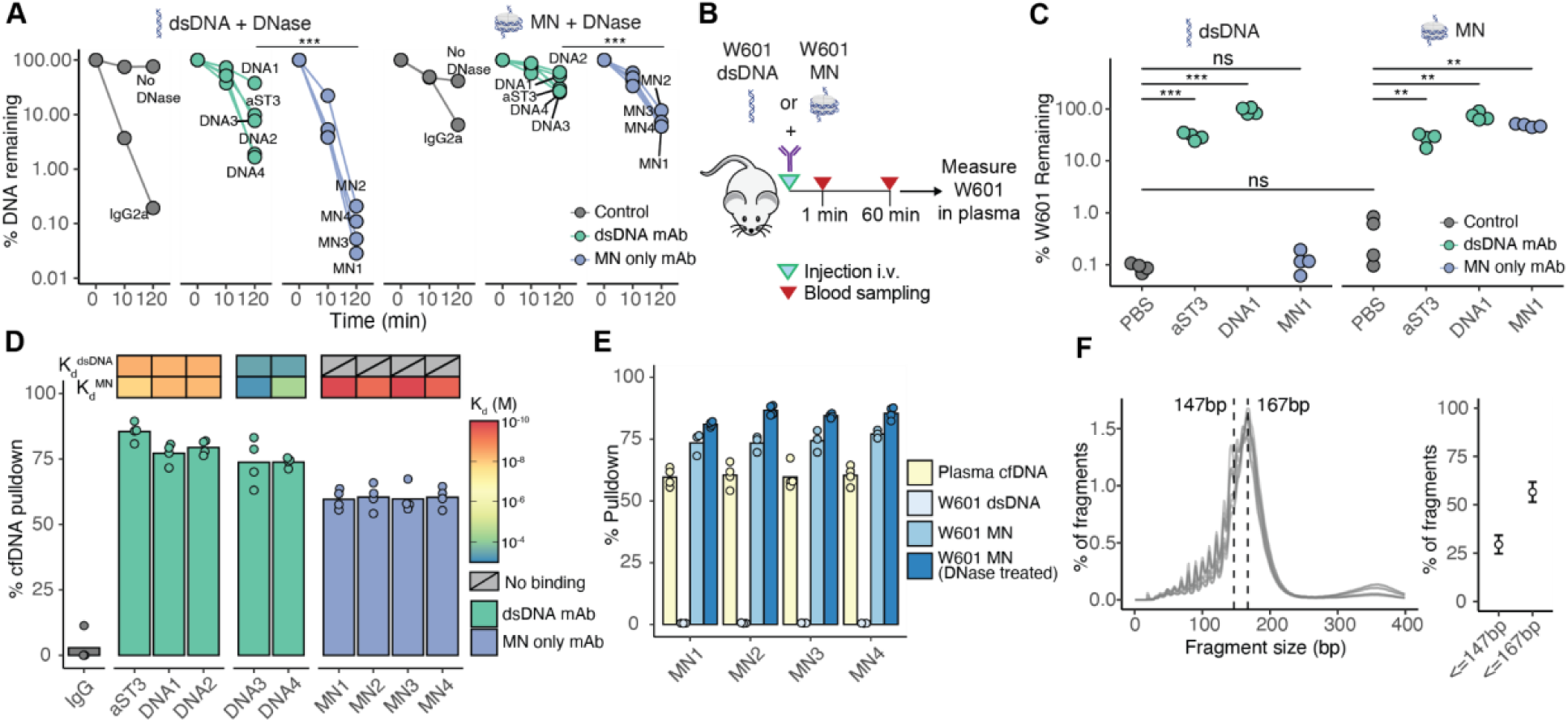
Impact of engineered mAb priming agents on cfDNA clearance and binding cfDNA in plasma. A) Degradation of free dsDNA and dsDNA in mononucleosomes in presence of 0.2U/mL of DNase I and mAb priming agents. IgG2a is an unrelated control antibody. Each dot represents the mean of two replicates. B) Experimental approach for testing impact of priming agents on clearance of free dsDNA and MN. C) Percentage of W601 DNA remaining in plasma 1 hour after injection of W601 either in free dsDNA form (left) or as MN (right), with or without dsDNA binding and MN binding priming agents. D) Percentage of cfDNA isolated from mouse plasma by various priming agents via immunoprecipitation using mAb-coupled magnetic beads, adjusted for background binding to beads alone. E) Percentage of cfDNA, W601 free dsDNA, W601 MN, and W601 MN with mild DNase I treatment isolated from mouse plasma using each MN-only mAb priming agent. F) cfDNA fragment length distribution in mice (n=8) with dashed lines at 167bp and 147bp (left) and percent of fragments <=147 bp and <=167 bp (right). * p < 0.05, ** p < 0.01, *** p < 0.001, ns - not significant.

Cell-free DNA uptake by organs, such as the liver and kidneys, is also a major clearance pathway for cfDNA.^18^ Given the impact of the form of cfDNA (free dsDNA versus MN) and specificity of priming agents (dsDNA versus MN) on nuclease-mediated degradation of DNA *in vitro*, we next asked how these variables affect the clearance of cfDNA in circulating plasma *in vivo*, where organ uptake also contributed to the clearance of cfDNA. We injected either free dsDNA or MNs into BALB/c mice with or without mAb priming agents and measured levels of the injected DNA in plasma via serial blood draws (**Fig. 2B**). The known Widom601 sequence (W601) was used in the injected preparation and its level in plasma was quantified by quantitative PCR (qPCR).^9,24^ Free dsDNA and intact MNs were both rapidly cleared from plasma (**Fig. 2C)**, with a median of 0.09% (range 0.07%-0.11%) of free dsDNA and 0.38% (range 0.1%-0.8%) of MN-bound dsDNA remaining 60 minutes after injection (p=0.10). In contrast to degradation by nucleases *in vitro*, clearance of free and MN-bound dsDNA from plasma *in vivo* was similarly rapid, suggesting that nuclease degradation may be a minor contributor to clearance of MN from plasma *in vivo*.

To investigate the impact of these mAbs on clearance kinetics of free and MN-bound dsDNA, we tested aST3 (K_d_^dsDNA^ 4.6nM, K_d_^MN^ 15.3nM); DNA1, dsDNA-binding mAb with high avidity to dsDNA (K_d_^dsDNA^ 5.2nM, K_d_^MN^ 5.9nM); and MN1, a mAb that only binds MN (K_d_^MN^ 0.14 nM and no binding to dsDNA). Both aST3 and DNA1 significantly reduced the clearance of free dsDNA: median 30% remaining at 60 min for aST3 and median 93% remaining at 60 min for DNA1. In contrast, MN1 had no impact on clearance of free dsDNA at 60 minutes (median 0.12% remaining), consistent with MN1 not binding to free dsDNA (**Fig. 2C, left; Fig. S2A)**. MN1 significantly reduced the clearance of MN-bound dsDNA (median 48% remaining at 60 min), as did aST3 and DNA1 (median 29% and 70% remaining at 60 min, respectively), consistent with the ability of the three mAbs to bind mononucleosomes (**Fig. 2C, right; Fig. S2A**). MN2 – another mAb with high avidity to MN only (K_d_^MN^ 0.44 nM, no binding detected to dsDNA) – showed similar results (**Fig. S2A,B)**. Together, these data suggest that both dsDNA and MN-only binding mAbs can slow the clearance of MN from plasma by virtue of high-avidity binding to MN. Only dsDNA binding mAbs, however, can slow the clearance of free dsDNA from plasma. Furthermore, although mAbs binding only MNs had no effect on DNase-mediated degradation of free and MN dsDNA *in vitro*, they significantly reduced the clearance of MN *in vivo*, consistent with the notion that degradation by nucleases may be a minor contributor to clearance of MN from plasma *in vivo* and suggesting that mAbs specific to MN could serve as priming agents for the portion of cfDNA that circulates as intact MN.

### dsDNA binders bind and isolate a higher fraction of endogenous cfDNA from plasma

Since physical interaction between mAbs and free dsDNA or intact MN is critical to their ability to inhibit degradation and clearance of cfDNA *in vitro* and *in vivo* (**Fig. 2A-C)**, we next sought to characterize the interaction of our mAbs with endogenous cfDNA. We performed immunoprecipitation of cfDNA from plasma from both BALB/c mice and humans using mAbs pre-coupled to magnetic beads, and estimated the fraction of cfDNA isolated. All mAbs isolated endogenous cfDNA from murine plasma compared to control, confirming that both intact mononucleosomes and dsDNA can serve as targets for priming agents (**Fig. 2D, Materials and Methods**). Similar results were observed with human plasma (**Fig. S2C**). Antibodies with both strong and weak avidities for dsDNA isolated the majority of cfDNA from plasma, with the stronger binders isolating only slightly more cfDNA compared to weaker binders despite the >1000x difference in K_d_ (**Fig. 2D**, **Fig. 1D**). This likely reflects the avidity boost from immobilization of mAbs in close proximity on the bead surface. Consistent with this result, when we used a different immunoprecipitation method allowing for endogenous IgGs to compete with our mAbs for binding to beads, the pulldown efficiency dropped significantly for the weak dsDNA binders but was maintained for strong dsDNA binders (**Fig. S2D**). Antibodies binding MN only isolated a lower fraction of cfDNA from plasma compared to the dsDNA specific binders, despite having the highest avidities to MN in our panel (**Fig 1D**). As cfDNA is a heterogeneous mixture of MN, partially disassembled MN, and free fragments of dsDNA, we reasoned that the lower efficiency of recovery for mAbs binding MN only may be partly due to their high specificity for intact MN, leaving free dsDNA and partially disassembled MN unbound. We examined the recovery of spiked-in W601 free dsDNA, W601 MN, and W601 MN after mild treatment with DNase 1 to reduce any free dsDNA component in the MN mixture. None of the MN-only binders could isolate W601 free dsDNA from plasma, consistent with the interferometry recording showing no binding to free dsDNA (**Fig. 2E**, **Fig. 1C left**). In contrast, MN-only binders isolated ∼75% of spiked in W601 MN (versus ∼60% of cfDNA), and the fraction of isolated W601 increased slightly after mild treatment with DNase I to reduce W601 free dsDNA in the MN preparation (**Fig. 2E**). Although cfDNA has a modal fragment length of 167-bp consistent with the length of dsDNA wrapped around histones with linker tails, we found that ∼30% of fragments in murine plasma had a length less than the 147-bp length associated with a nucleosome core particle, and may represent sub-nucleosomal or free fragments of dsDNA not bound by MN-only binders (**Fig. 2F**), consistent with the results from immunoprecipitation studies. Together, these data suggest that MN binders may not be able to bind the fraction of cfDNA that exists as partially-disassembled MN or free dsDNA, which could negatively affect their performance as priming agents.

### cfDNA binding mAbs increase recovery of ctDNA in tumor-bearing mice

The characterization of our different cfDNA-binding mAbs revealed a diversity of specificities, interaction kinetics, and avidity to cfDNA, and these traits, in turn, correlated with binding to cfDNA and protection from degradation and clearance *in vivo*. These analyses may not fully recapitulate the biology of endogenous cfDNA in plasma *in vivo*, however. To determine whether our mAbs could act as priming agents and whether specificity matters for priming, we next sought to evaluate the impact of our mAb priming agents on endogenously-released cfDNA and ctDNA *in vivo*.

We tested all of the mAbs that showed any binding to dsDNA or MN in tumor-bearing mice (**Fig. 3A**). As in prior work, we used a dose of 4mg/kg and collected plasma both before administration and 2 hours afterwards for each agent in BALB/c mice harboring murine colon carcinoma MC26 lung tumors. We measured ctDNA using a next-generation sequencing (NGS)-based assay with a custom panel tracking 1,822 SNVs in MC26, and measured the “priming effect” as the change in the number of ctDNA molecules per mL of plasma in each mouse after administration of a priming agent. Total cfDNA and ctDNA increased in mice after administration of mAb priming agents (**Fig. 3B-C**), with similar tumor fractions compared to PBS (control) (**Fig. S3A**). The magnitude of the priming effect varied considerably. The mAbs with high avidity for dsDNA (aST3, DNA1, DNA2) resulted in the largest increase in amounts of ctDNA, whereas weak dsDNA binders (DNA3, DNA4) had little to no effect. Surprisingly, although MN-only binders had the lowest K_d_^MN^ values in our panel (all <1 nM), they all had a modest priming effect, resulting in 2.4-3.2x increase in ctDNA, as compared to 12-13x for the best dsDNA binders. Consistent with these results, priming agents resulted in detection of a larger percentage of mutations from the 1,822-SNV panel compared to no priming, with the greatest improvements observed for high-avidity dsDNA binders (**Fig. S3B**).

**Figure 3:**
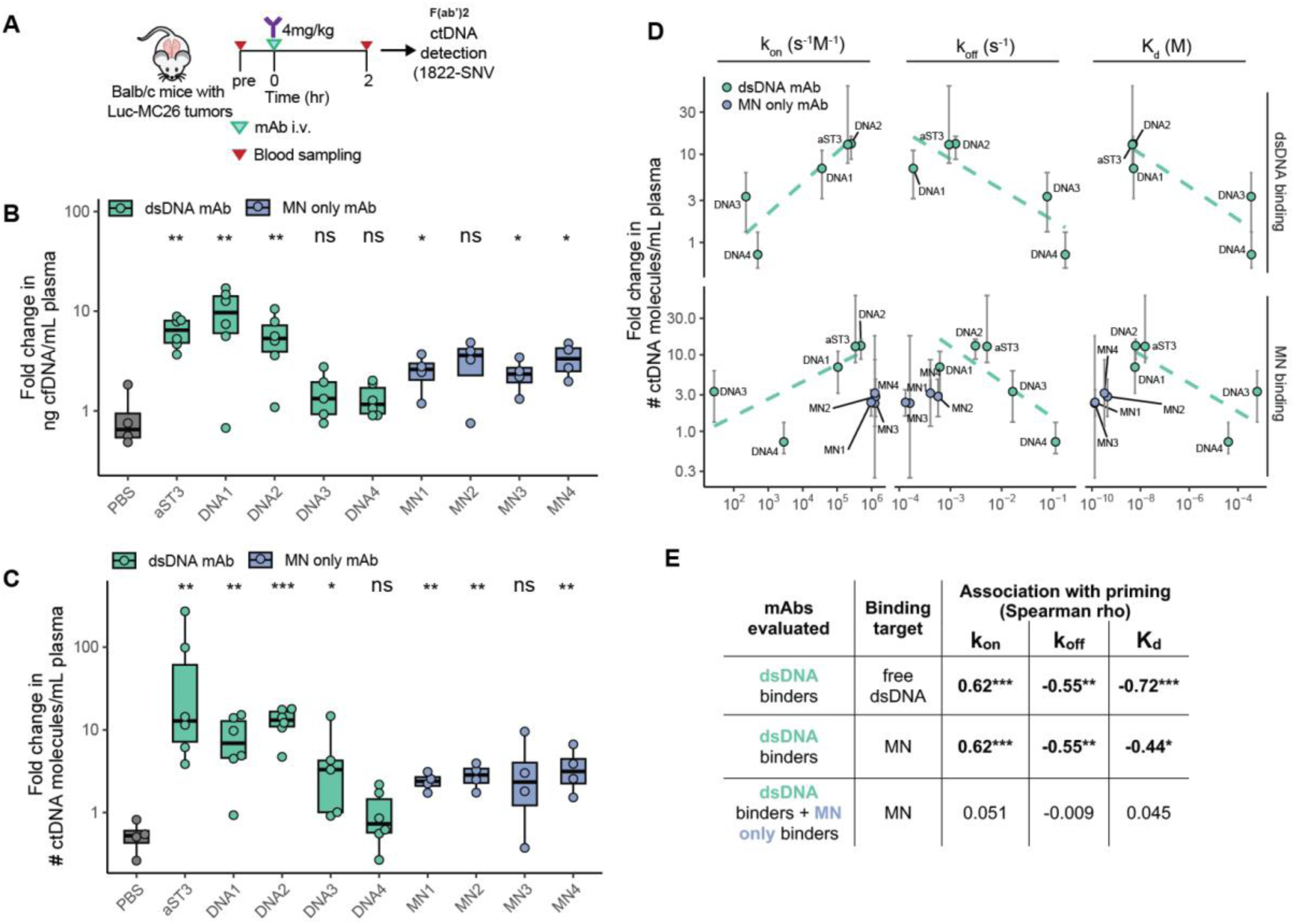
Engineered mAb priming agents increase ctDNA recovery in vivo. A) Experimental schematic for testing impact of priming agents on cfDNA and ctDNA recovery. Priming agents were injected at 4mg/kg IV, and blood was collected before and 2 hours after injection. Total cfDNA and ctDNA levels were compared before and after administration of priming agents in each mouse to measure fold change. B) Fold-change in plasma cfDNA concentration after administration of priming agents. C) Fold-change in number of ctDNA molecules detected in plasma after administration of priming agents. D) Correlation of priming effect, as measured by change in number of ctDNA molecules detected in plasma, and interaction kinetics to dsDNA or MN. E) Spearman correlation coefficients for associations in panel D. * p < 0.05, ** p < 0.01, *** p < 0.001, ns - not significant. Box plots represent median and interquartile range.

We examined the relationship between binding kinetics and the magnitude of the observed priming effect. For dsDNA binders, the priming effect was correlated with k_on_, k_off_, and K_d_ of the priming agent to dsDNA and MN (**Fig. 3D,E**), with faster k_on_, slower k_off_, and lower K_d_ associated with greater priming effect. This relationship between avidity and priming did not hold when we analyzed dsDNA and MN-only binders in aggregate; MN-only binders underperformed as priming agents relative to the strength of their binding interaction with MN (**Fig. 3D, 3E bottom row**). Together, these findings indicate dsDNA is the key target for antibody-based priming agents, with K_d_^dsDNA^ < 10nM associated with the best priming effects. MN-only binders, despite their remarkably high avidity to MN (K_d_^MN^ < 1nM) and ability to delay MN clearance from plasma (**Fig. 2C**), did not perform well as priming agents.

### Impact of stabilization of cfDNA by mAbs in plasma

Since mAb priming agents directly bind cfDNA and prolong its half-life in plasma, we next asked what impact this interaction had on fragmentation of cfDNA and the representation of the genome in cfDNA. We reasoned that if cfDNA persists in the bloodstream, it may be exposed to extracellular nucleases for longer, and that this may alter its fragmentation if the priming agent does not also confer sufficient protection from nucleases. It is further possible that priming agents with superior protection from nucleases may enhance the recovery of short fragments of cfDNA not protected by histone proteins (**Fig. 2A**) which may otherwise be underrepresented or lost due to nuclease-mediated digestion. To investigate these questions, we performed low coverage (1-4x) whole genome sequencing of plasma from mice following administration of DNA-binding mAb priming agents. Mice receiving PBS had the expected size distribution of cfDNA with a modal fragment size of 167 bp (**Fig. 4A)**. Several dsDNA-binding priming agents recovered a higher fraction of short fragments <= 120bp than PBS control (**Fig. 4A,B)**. Higher amounts of short DNA were seen with mAbs that bound dsDNA with high avidity (aST3, DNA1, and DNA2), whereas those that bound dsDNA weakly (DNA3, DNA4) had similar size distributions to controls. MN binders (MN1-MN4) were associated with a distribution of fragment-length with a more prominent mononucleosomal peak consistent with preferential protection of intact MN in plasma (**Fig 4A**). Despite having nearly identical K_d_^dsDNA^ (**Fig. 1D**), aST3, DNA1, and DNA2 were associated with different amounts of short fragments of cfDNA (**Fig. 4A,B**). The abundance of short fragments of cfDNA was associated with faster dissociation rates (k_off_) from dsDNA (**Fig. 4C**) and was inversely correlated with extent of protection from DNase *in vitro* (**Fig. 4D**). The lowest abundance of short cfDNA was observed for the priming agent DNA1 which yielded the least degradation by nucleases. This effect may be due to more stable interactions with dsDNA as reflected by its slow k_off_, which may be an important parameter regarding its nuclease protection ability.

**Figure 4:**
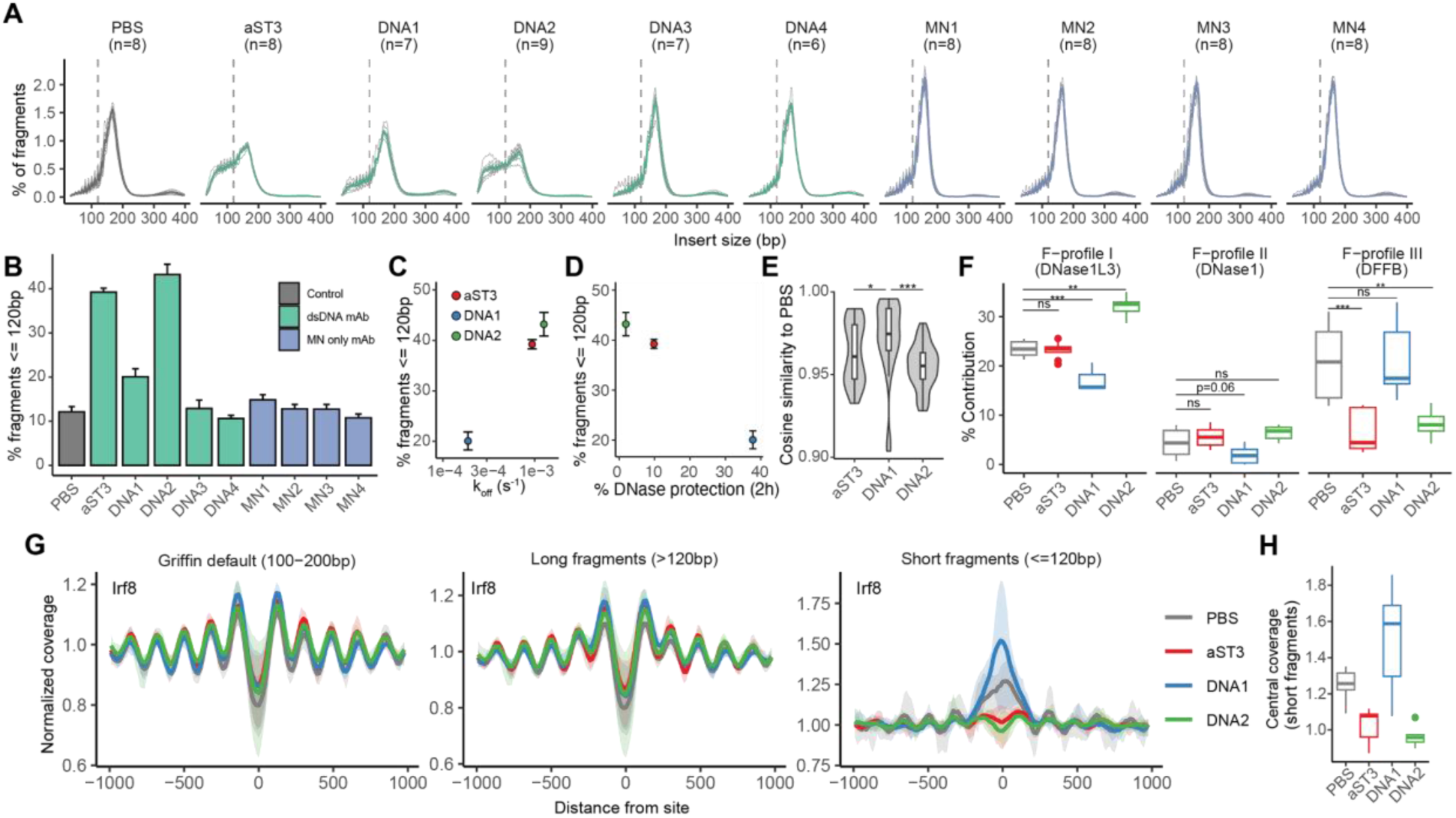
cfDNA fragmentation and end-motifs after administration of priming agents. A) Plasma cfDNA fragment-size distribution in mice following administration of priming agents. The colored line in each panel is the average across all mice in the group and gray traces are distributions for individual mice in the group. Dashed vertical line at 120-bp. B) Percent of short fragments (<=120bp) in cfDNA with each priming agent. C) Correlation of dissociation (k_off_) rate with dsDNA with abundance of short cfDNA fragments in plasma after priming agent administration. D) Correlation of DNase protection, as measured by % dsDNA remaining at 2 hours, and abundance of short cfDNA fragments in plasma. E) Cosine similarity of 4-mer end-motif profiles for plasma samples after priming agent administration versus PBS administration. F) Contribution of F-profiles I-III, associated with DNase1L3, DNase1 and DFFB activity respectively, to 4-mer end-motifs in cfDNA after administration of PBS or priming agents. G) Coverage patterns around Irf8 binding sites using Griffin’s default fragment filter (100-200bp), long fragments (>120bp, range 121-500bp), and short fragments (<= 120bp, range 15-120bp). Dark lines represent mean normalized coverage for each priming agent and bands indicate the range across samples for each agent. H) Central coverage at Irf8 sites by short fragments (<= 120bp, range 15-120bp). * p < 0.05, ** p < 0.01, *** p < 0.001, ns - not significant. Box plots represent median and interquartile range.

Given the impact of aST3, DNA1, and DNA2 on cfDNA fragmentation and differences in distribution of fragment length (despite similar K_d_^dsDNA^ and similar priming effect), we further analyzed end-motif patterns in cfDNA after administration of these three agents. Sequences at the end of fragments of cfDNA (end-motifs) are non-random and can reflect the activity of specific nucleases.^20,25,26^ To analyze cfDNA end-motifs after using priming agents, we calculated the frequency of each of 256 possible 4-mers, defined as the terminal four nucleotides at the 5’ end of each fragment, in cfDNA samples after administration of aST3, DNA1, DNA2, or PBS (control). Additional filters were incorporated to account for variable length unique molecular identifiers (UMIs) in our adapters and the raw frequencies were adjusted based on the frequency of 4-mer’s in the mouse genome, consistent with other studies (**Materials and Methods**).^26^ The end-motif profile of DNA stabilized by DNA1 most closely resembled that of samples without priming agents (PBS only) (**Fig. 4E**, **Fig. S4A**), while the profiles of samples from mice treated with aST3 and DNA2 were most similar to one another, compared to samples from mice treated with either agent versus DNA1 (**Fig. S4B).** To determine whether subsets of end-motifs were differentially represented in cfDNA after priming, we compared the frequency of end-motifs in samples from mice given aST3, DNA1, or DNA2 priming agents versus control samples (PBS treated) and identified 24 end-motifs that were enriched and 33 end-motifs that were depleted among the three comparisons, with the majority in samples from mice given aST3 and DNA2 samples (**Fig. S4C,E, Table S4**). None were enriched or depleted in samples from mice receiving IgG2a control (**Fig. S4D**). Almost all enriched end-motifs had C-ends or T-ends, and the majority of depleted end-motifs had A-ends and a minority had G-ends (**Fig. S4E, Table S4**), with significant overlap in enriched (p = 4.0×10^-7^, Fisher’s exact test) and depleted (p = 2.1×10^-11^, Fisher’s exact test) end-motifs between aST3 and DNA2 (**Fig. S4F**). Only six end-motifs were enriched after DNA1 and none were depleted, suggesting that endogenous end-motifs are better preserved with DNA1 (**Fig. S4E**).

Activity of major nucleases involved in cfDNA fragmentation (DNase1, DNase1L3, DFFB) results in stereotypic fragment end-motif sequences.^20,25,27,28^ We hypothesized that changes in cfDNA end-motifs after priming agents could reflect changes in the relative contribution of various nucleases to the fragmentation of cfDNA. To test this hypothesis, we used non-negative least squares to deconvolve the end-motif profile in each sample into the end-motif F-profiles that describe the distinct patterns of end-motif cleavage found in cfDNA.^26^ F-profiles I, II and III reflect the end-motif patterns associated with DNase1L3, DNase1 and DFFB activity, respectively.^26^ We found that F-profile III (intracellular nuclease DFFB) contributed significantly less in samples from mice treated with aST3 and DNA2 compared to control (PBS only) and DNA1. These data are consistent with depletion of A-end fragments with these agents (**Fig. 4F, Fig. S4G**), and DNA2 was associated with a higher contribution of F-profile I (DNase1L3). DNA1, in contrast to aST3 and DNA2, had a similar level of contribution from F-profile III (DFFB) as control samples (PBS only), with lower contributions from F-profile I (DNase1L3) and F-profile II (DNase1) (**Fig. 4F**). The lower contribution of extracellular nucleases (DNase1L3 and DNase1) in samples from mice receiving DNA1 *in vivo*, potentially less than PBS treated samples, would be consistent with its superior protection of dsDNA from nuclease digestion compared to those with aST3 and DNA2 *in vitro* (**Fig 4D**), and may reflect better protection from extracellular nucleases DNase1 and DNase1L3. This result raises the question of whether or not additional recovery of short fragments of cfDNA fragments by DNA1 as compared to PBS may reflect the enhanced recovery of short, biologically-informative fragments that are otherwise lost to nuclease digestion without a priming agent.

To better understand the origins of short fragments of DNA recovered from mice treated with aST3, DNA1, and DNA2, we next examined patterns of nucleosomal and subnucleosomal coverage with Griffin.^29^ Using Griffin’s default settings for fragment length (100-200bp), we observed highly similar periodic coverage fluctuations characteristic of nucleosomal protection around binding sites of a blood-specific transcription factor, Irf8 (**Fig. 4G, left**), suggesting that priming agents generally preserve the nucleosomal patterns in cfDNA. Given the size distributions observed in our samples, we next analyzed coverage patterns by long (>120bp, range 121-500bp) and short (<=120bp, range 15-120bp) fragments. Long fragments of cfDNA correspond generally to regions of nucleosome and genome coverage by these fragments fluctuates by nucleosome occupancy, whereas short fragments of cfDNA correspond to regions between nucleosomes and coverage by short fragments is higher at transcription factor binding sites.^30^ After administration of priming agents or control PBS, we observed a similar pattern of nucleosomal coverage in long fragments (>120bp), consistent with the nucleosomal origin of these fragments (**Fig. 4G, middle**). Among short fragments (<=120bp), we observed an expected rise in coverage at the binding site (“central coverage”) compared to controls (PBS). Among priming agents, DNA1 administration was also associated with high central coverage, with a peak that was more pronounced compared to PBS control. In contrast, no central coverage peak was observed with treatment of aST3 or DNA2 (**Fig. 4G, right; Fig. 4H**). Similar patterns were observed at binding sites of other major blood transcription factors Spi1 and Lyl1 (**Fig. S5A**). Together, these findings suggest that patterns of nucleosomal coverage among long fragments (>120bp) are unchanged with priming agents, consistent with the nucleosomal origin of these fragments. The excess of short fragments observed with aST3 and DNA2 likely originates from longer nucleosomal fragments that are digested into shorter fragments. In contrast, DNA1 is associated with a cfDNA fragmentation and coverage profile that is more consistent with PBS control. Notably, DNA1 was associated with higher central coverage by short fragments compared to PBS control across a range of transcription factors (**Fig. S5B**), suggesting that the excess of short cfDNA observed with DNA1 may reflect biologically-informative short fragments of cfDNA that may otherwise be rapidly cleared and thus, lost from plasma, in the absence of a priming agent. These features would be advantageous in a priming agent for better preserving informative features of cfDNA.

### The Fc domain is dispensable to priming activity

The Fc domain mediates the effector function and pharmacokinetics (PK) of mAbs through interactions with Fcɣ receptors and the neonatal Fc receptor (FcRn).^31,32^ We previously showed that abrogating interaction of the Fc domain with Fcɣ receptors is critical to the priming effect.^9^ Whether the presence of the Fc domain is strictly necessary for the priming effect remains unknown. We tested the role of the Fc domain by comparing aST3 to a F(ab’)2 with the variable domain of aST3. The F(ab’)2 was generated via IdeZ digestion of aST3 without the L234A/L235A/P329G mutations, which confer IdeZ resistance. As IdeZ digests below the IgG2a hinge region, residues L234 and L235 were retained in the F(ab’)2 fragment. Although the Fc domain has been reported to be required for dsDNA binding of some anti-DNA mAbs,^17^ we found that the F(ab’)2 bound dsDNA and MN with similar avidity as aST3 (K_d_^dsDNA^ = 7.6nM, K_d_^MN^ = 7.4nM), suggesting that the variable domain of aST3 alone is sufficient for binding to dsDNA and MN with high avidity (**Fig. S6A,B**). To investigate the pharmacokinetics of F(ab)’2 compared to aST3, we labeled F(ab’)2, aST3, and an unrelated IgG2a mAb with the L234A/L235A/P329G mutations with Cy7 and measured mAb concentrations in plasma over time (**Fig. 5A**). Compared to the IgG2a mAb, both aST3 and F(ab’)2 were cleared from plasma more rapidly (**Fig. 5B**). This result suggests that the variable domain of antibodies could play a role in plasma pharmacokinetics, and specifically, that binding to cfDNA results in more rapid clearance from plasma. Furthermore, F(ab’)2 was cleared more rapidly than aST3, which is expected given its smaller size and lack of Fc domain (**Fig. 5B**). Plasma samples collected 2 hours post priming agent administration revealed that the majority (>50%) of the injected dose of each agent was still present in plasma, however, the concentration of F(ab’)2 was slightly lower than that of aST3 (58% +/- 3.6% and 72% +/- 1.8% remaining, respectively), and both were lower than the concentration of IgG2a control (92% +/- 1.9% remaining) (**Fig. 5C**).

**Figure 5:**
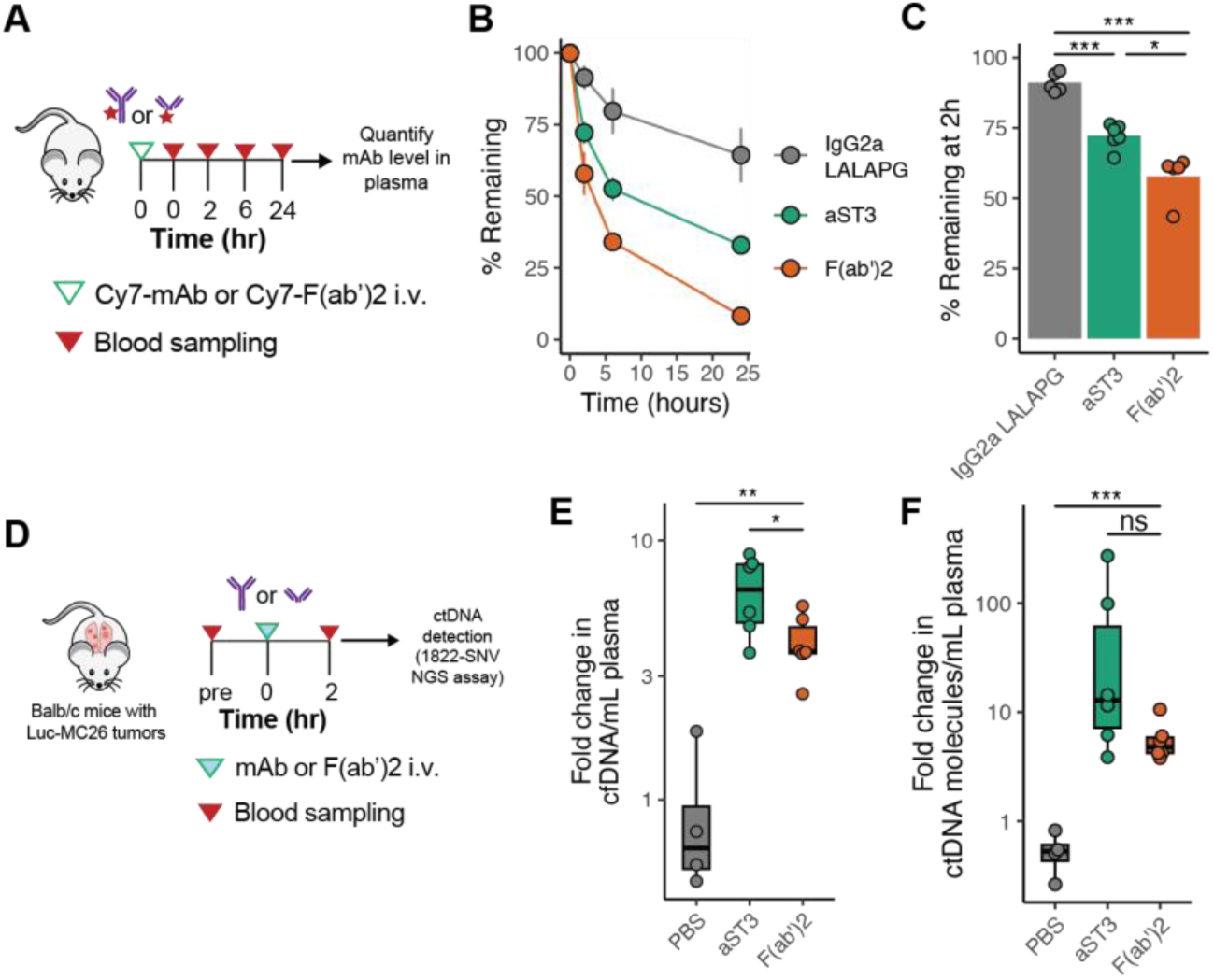
Impact of Fc domain on priming. A) Experimental approach for measuring pharmacokinetics of aST3, F(ab’)2, and IgG2a LALAPG in plasma. Fluorophore-labeled full-length mAbs (aST3 or IgG2a LALAPG isotype control) were injected IV and blood was sampled at 1min, 2h, 6h, and 24h after injection. All values were normalized to the 1min plasma levels to measure percent remaining. B) Percent of injected agents remaining in plasma over time. C) Percent of agent remaining in plasma at 2 hours. D) Experimental approach for testing priming effect of F(ab’)2 and aST3 on cfDNA and ctDNA recovery. E) Fold-change in plasma cfDNA concentration after administration of each priming agent. F) Fold-change in number of ctDNA molecules detected in plasma after administration of priming agents. * p < 0.05, ** p < 0.01, *** p < 0.001, ns - not significant. Points represent mean and intervals represent 95% confidence intervals in B. Box plots represent median and interquartile range.

Given that >50% of the injected dose of F(ab’)2 remained in plasma after 2 hours, we hypothesized that it may still elicit a priming effect despite its more rapid clearance compared to aST3. We tested the priming effect of F(ab’)2 at a dose of 2.67 mg/kg (the same molar dose as intact mAb at 4mg/kg). F(ab’)2 was active as a priming agent, resulting in more total cfDNA and more ctDNA in plasma compared to control PBS (**Fig. 5D-F**), suggesting that the Fc domain is dispensable to the priming effect. The observed magnitude of priming was numerically slightly lower compared to aST3 in recovery of both cfDNA and ctDNA (**Fig. 5E-F**) although the difference did not reach statistical significance for recovery of ctDNA (**Fig. 5F**). Overall, these results demonstrate that the priming effect does not require an intact Fc domain, and the plasma concentrations of a F(ab’)2 are sufficient after 2 hours to elicit a priming effect, despite its more rapid clearance. Further optimization of the administered dose and sequence of the F(ab’)2 may be required, however, to further enhance the observed priming effect to match or exceed that which is observed for full-length mAbs.

### Rationally-designed single-chain dsDNA binders as priming agents

Our investigation of mAbs as priming agents identified dsDNA, rather than MN, as the key binding target, and suggested that stronger binding to dsDNA, characterized by faster k_on_, slower k_off_, and lower K_d_, was associated with an improved priming effect. We next asked whether these insights into mAbs could be used to guide the design of new priming agents. We reasoned that if binding to dsDNA is the key property for priming cfDNA, then a non-immunoglobulin dsDNA binding protein could similarly elicit a priming effect.

We designed novel candidates for priming agents using sso7d, a 7 kDa dsDNA-binding protein found in the thermophilic archaeon *Sulfolobus solfataricus*.^33^ Sso7d has been investigated as a binding scaffold due to its high thermal stability, tolerance of pH fluctuations, lack of cysteines and glycosylation sites, and ease of expression.^34,35^ It has been fused to DNA polymerases to enhance processivity, and agents using the sso7d backbone are being explored in applications for affinity-based purification, and for diagnostic and therapeutic agents.^36–38^ We generated single-chain, sso7d-based agents by fusing either one (“mono-sso7d”), two (“bi-sso7d”) or four (“tetra-sso7d”) domains of sso7d to a murine IgG2a-LALAPG Fc domain (**Fig. 6A**). Long [(G4S)_5_] and short [(G4S)_3_] flexible linkers were used for sso7d-Fc and tandem sso7d fusions, respectively, and knob-in-hole mutations^39^ were added to the Fc domain of mono-sso7d to promote heterodimerization (**Table S1**).

**Figure 6:**
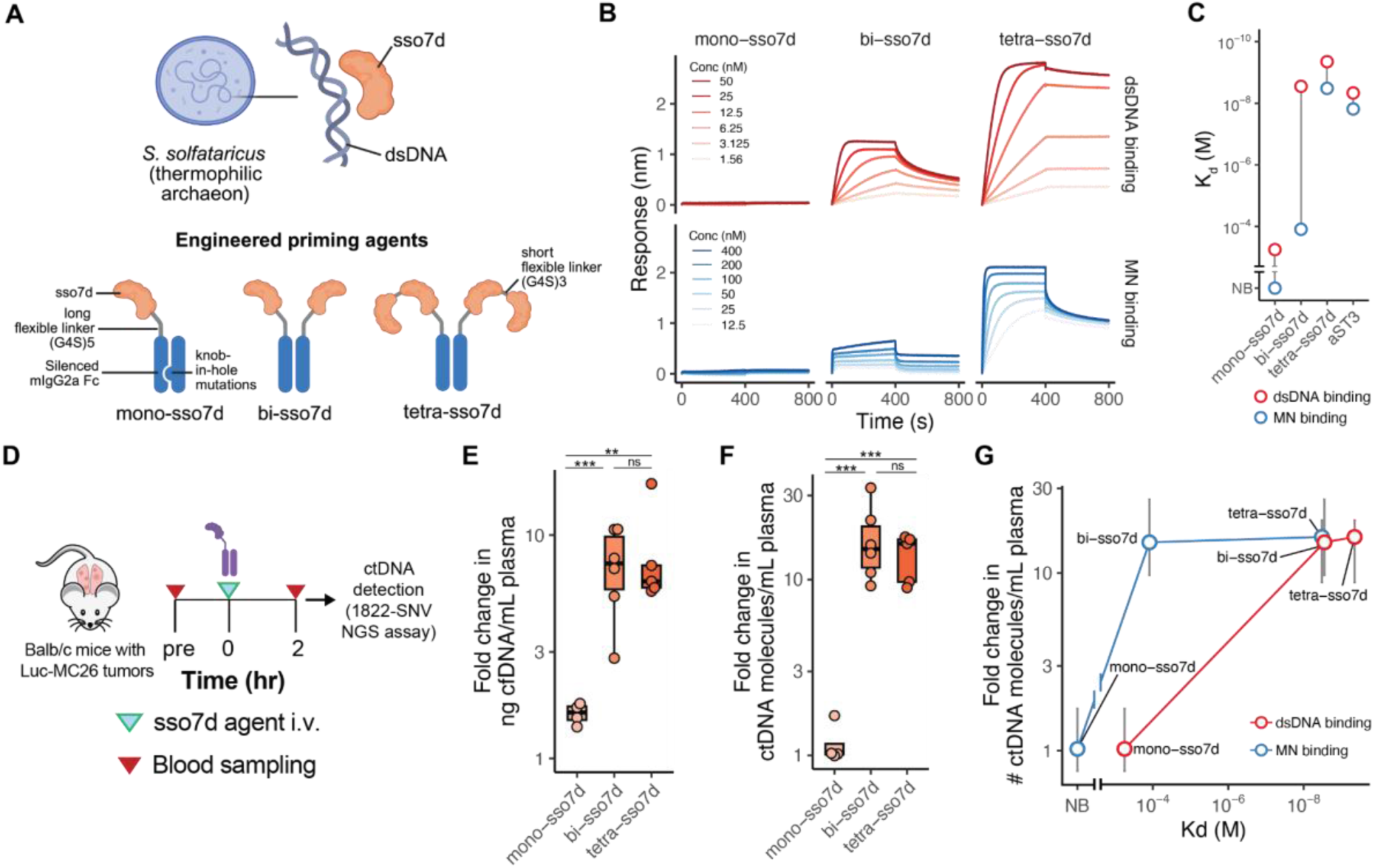
Rationally-designed single-chain dsDNA binders improve ctDNA recovery. A) Design of priming agents using the 7 kDa dsDNA binding protein sso7d. Mono-valent, bi-valent, and tetra-valent constructs were designed using flexible short (G4S)_3_ and long (G4S)_5_ linkers fused to murine IgG2a CH2-CH3 domain carrying the LALAPG silencing domain, with addition on knob-in-hole mutations to promote heterodimerization for the mono-sso7d construct. B) Interferometry recordings of the interaction of sso7d priming agents with free dsDNA (“dsDNA binding”) and MN (“MN binding”). For binding to dsDNA, the range of priming agent concentrations tested was 1.56-50nM, whereas for binding to MN, the range of concentrations tested was 12.5-400nM in order to capture weaker interactions. C) Estimated avidity (K_d_) for binding to dsDNA and MN for sso7d priming agents compared to aST3. D) Experimental approach for testing the effect of sso7d priming agents on cfDNA and ctDNA recovery. E) Fold-change in plasma cfDNA concentration after administration of each priming agent, based on qPCR quantification. F) Fold-change in number of ctDNA molecules detected in plasma after administration of priming agents. G) Priming effect versus K_d_ to dsDNA and MN for sso7d priming agents. * p < 0.05, ** p < 0.01, *** p < 0.001, ns - not significant. Points represent mean and intervals represent 95% confidence intervals in G. Box plots represent median and interquartile range.

We first measured the binding kinetics of the sso7d constructs with free dsDNA and MN using BLI **(Fig. 6B-C, Table S3**). Mono-sso7d showed no detectable binding to MN and very weak binding to free dsDNA (K_d_^dsDNA^ 562 μM), detectable only at high concentrations of the molecule (**Fig. S6C**). In contrast, bi-sso7d showed much stronger binding to free dsDNA (K_d_^dsDNA^ 2.8 nM) and detectable, albeit weak, binding to MN (K_d_^MN^ 123 μM), highlighting the critical role of monogamous bivalency and avidity effects in enabling binding of dsDNA-targeted priming agents to dsDNA. Addition of more sso7d domains in tetra-sso7d resulted in slightly stronger binding to free dsDNA (K_d_ ^dsDNA^ 0.45 nM) and significantly stronger binding to MN (K_d_ ^MN^ 3.3 nM) compared to bi-sso7d.

We next evaluated the priming effect of the sso7d-based agents in tumor-bearing mice (**Fig. 6D**). Mono-sso7d did not result in changes in recovered yields of total cfDNA or ctDNA, likely reflecting its weak interaction with free dsDNA and MN (**Fig. 6E-F**). By contrast, bi-sso7d and tetra-sso7d both led to increased yields of endogenous cfDNA and ctDNA, resulting in median 14.9-fold and 16-fold increases in ctDNA (**Fig. 6F**). Furthermore, the stronger binding to free dsDNA in bi-sso7d compared to mono-sso7d was associated with a better priming effect, whereas stronger binding to MN in tetra-sso7d (compared to bi-sso7d) did not further improve the priming effect (**Fig. 6G**). This discrepancy between the impact of binding to dsDNA versus MN further emphasizes dsDNA as the key target for priming agents, and suggests that MN binding alone is neither sufficient nor necessary for priming. Together, these results demonstrate that the factors associated with the priming effect of antibody-based priming agents can guide the design of new molecules with the ability to act as priming agents for liquid biopsies.

## Discussion

In this study, we showed that multiple mAbs that bind cfDNA can have a priming effect that increases the recovery of ctDNA from a blood draw. Through biochemical characterization and correlation with *in vivo* priming experiments, we identified the features associated with this priming effect. *In vitro*, dsDNA binders provided better protection from nucleases and bound a larger fraction of cfDNA compared to MN binders. *In vivo*, antibodies that bound dsDNA directly with fast association, slow dissociation, and low overall K_d_ performed best as priming agents. Antibodies that bound MNs exclusively, despite having some of the lowest K_d_’s measured in this study (< 1nM), did not perform as well as dsDNA binders as priming agents. Furthermore, the Fc domain was dispensable to priming. Using these principles, we designed novel single-chain priming agents using the specific dsDNA-binding domain sso7d, and demonstrated that bivalent and tetravalent agents both had high avidity to dsDNA and had a priming effect in tumor-bearing mice. Our work here expands the breadth of antibody-based priming agents, identifies the key determinants of their function, and shows the generalizability of these findings for designing new molecules as priming agents (**Fig. 7**).

**Figure 7:**
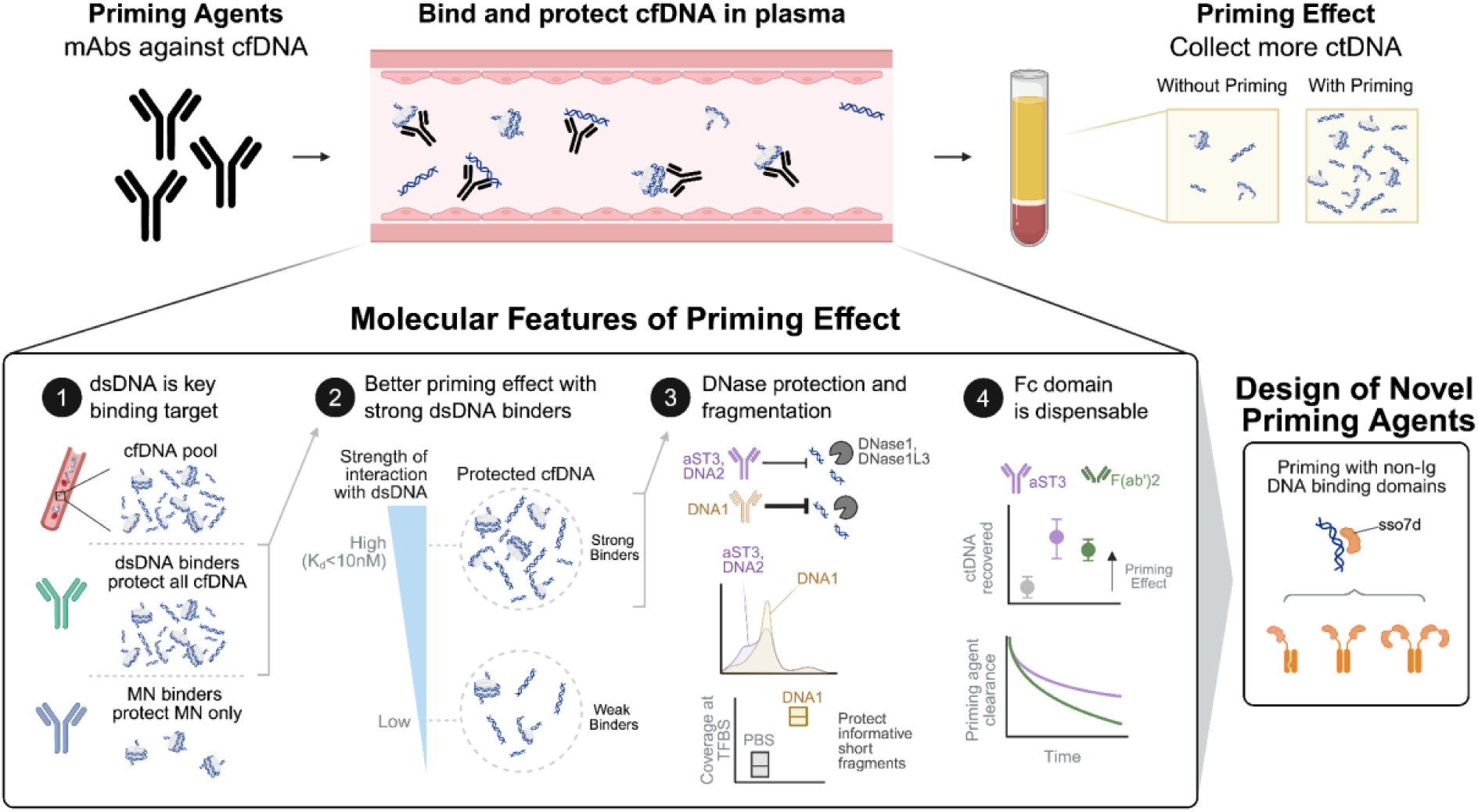
Molecular determinants of antibody-mediated cfDNA priming. Antibody-based priming agents bind cfDNA in the bloodstream and protect it from clearance, enabling more to be collected in a subsequent blood draw. This study identified the key molecular determinants of priming activity. The optimal target binder was dsDNA, rather than mononucleosomes, and the priming activity was correlated with strength of binding to dsDNA, with the best priming agents having K_d_^dsDNA^ < 10nM. The best dsDNA-binding priming agents had different magnitudes of DNase protection ability and impact on cfDNA fragmentation. In particular, one agent, DNA1, best preserved the endogenous fragmentation profile in cfDNA and protected short, informative cfDNA fragments at transcription factor binding sites from clearance. The Fc domain was found to be dispensable for the priming effect, suggesting that agents with more rapid clearance can still elicit a priming effect. Finally, we leveraged some of the principles identified in this study to engineer new single chain molecules that can similarly elicit a priming effect in tumor bearing mice, extending the space of priming agents to non-immunoglobulin dsDNA binding domains.

Our analysis of patterns of fragmentation, end-motif, and coverage following administration of aST3, DNA1, and DNA2 highlight other parameters that may be important in the selection of priming agents. We found that although the three agents had similar K_d_ ^dsDNA^ and K_d_^MN^, and similar priming effects, samples from mice treated with DNA1 resulted in a fragmentation profile more consistent with cfDNA from control samples, and was thus better able to preserve the baseline endogenous features of cfDNA. This property may be due, in part, to DNA1’s distinct interaction kinetic rates with dsDNA and its superior DNase-protection effect. All three agents preserved the nucleosomal coverage patterns in cfDNA by nucleosomal (>120bp) fragments. DNA1, however, yielded better preservation of short, biologically informative fragments of cfDNA associated with transcription factor binding sites, likely due to its superior ability to protect DNA from DNase degradation. For applications of priming agents in cfDNA-based liquid biopsies, agents such as DNA1 may offer advantages in the subset of assays that use features of fragmentation by better preserving the fragment and end-motif features of cfDNA while delaying its clearance and boosting its recovery, particularly of short fragments informative of transcription factor binding.

Our investigation of mAbs as priming agents also led to new insights into the biology of cfDNA. Short fragments of cfDNA have been observed in various conditions, including cancer and autoimmune diseases such as systemic lupus erythematosus (SLE).^40–42^ In SLE, including the subset caused by germline deficiency of DNase1L3,^42^ patients can develop endogenous anti-nuclear antibodies (ANAs) against dsDNA and chromatin. Fc-mediated interactions are key mediators of the autoimmune impact of ANAs,^43^ and our priming agents have been Fc-silenced for this reason. But whether the changes in cfDNA fragmentation observed in patients with cancer and autoimmune diseases are related to ANAs or due to other processes (e.g. changes in nuclease activity) remains unclear. In our data, a subset of mAb priming agents with weak protection from nucleases was associated with cfDNA fragment shortening in mice without autoimmune disease. Size shortening was observed with high avidity dsDNA binders and not seen with MN binders or weak dsDNA binders. Furthermore, the degree of size shortening was correlated with kinetics of interaction with dsDNA, and end-motif analysis highlighted the role of circulating nucleases and the degree of nuclease-protection from each priming agent in this process. Together, our observations suggest that a specific subset of ANAs that bind dsDNA with high avidity but with fast k_off_ and low ability to protect from DNase may be sufficient to cause size shortening of cfDNA. Since ANAs occur naturally in the population, including in many cancer patients, these results have important implications for methods that rely on cfDNA fragmentation for cancer detection,^44,45^ where the presence of anti-dsDNA antibodies should be considered when interpreting their results.

Antibody-based priming agents can boost sensitivity in applications where detection power is constrained by the amount of input material, such as MRD detection and tumor genomic profiling from plasma, as well as early cancer detection. By transiently inhibiting the pathways for biological clearance of cfDNA, antibody-based priming agents improve the amount of ctDNA collected in a blood draw and thus improve the detectable signal. Much work has been done to date on improving the performance of technologies *in vitro* and *in silico* to detect cfDNA. As a result, assays can now profile cfDNA samples to “saturation,” interrogating all fragments of cfDNA at up to thousands of loci genome-wide in search of molecules of ctDNA.^3–5^ Antibody-based priming agents could serve as a unique complementary technology to these advances, modulating the biology of cfDNA in the body to vastly improve the number of cfDNA molecules that can be profiled in a sample by these high-throughput, ultrasensitive methods. In this work, we showed the generalizability, determinants, and design principles for this approach, establishing the groundwork for systematic development of priming agents.

Our work raises several questions for future investigation. Although we investigated a panel of published sequences of antibodies against dsDNA and mononucleosomes, the space of potential binders against cfDNA is vast, and as our results with sso7d suggest, likely extends beyond immunoglobulin binding domains. Thorough investigation of this design space would benefit from strategies for high-throughput screening. Reliance on *in vivo* testing for priming effects remains a bottleneck for testing new priming agents and studying the biology of cfDNA. Development of *in vitro* and cell-based surrogates of the biology of cfDNA could help accelerate development and evaluation of priming agents. We examined the role of the Fc domain in performance of priming agents here, and found that despite more rapid clearance, F(ab’)2 fragments could still elicit a priming effect. More rapid clearance of priming agents would be a desirable property for clinical translation. Testing these strategies in a combinatorial fashion with different binding domains could yield further improvements for protein-based priming agents.

Patterns of nucleosome coverage were preserved in nucleosome-length fragments of cfDNA (100-200bp) after administration of priming agents. Strong dsDNA binder priming agents (aST3, DNA1, and DNA2) were associated with elevated levels of short fragments of cfDNA, to varying degrees. These changes may affect fragmentomics approaches that rely on the relative abundance of short versus long fragments of cfDNA. DNA1 best preserved the fragmentation properties of endogenous cfDNA and was associated with better recovery of short, biologically informative fragments of cfDNA at transcription factor binding sites, likely due to its superior ability to protect from DNase. These properties may be advantageous in a priming agent for cfDNA. We also note that because fragmentomics-based approaches do not profile cfDNA to saturation and typically use shallow sequencing, the fundamental constraint in their sensitivity is not total input mass but rather the relative abundance of ctDNA. Therefore, priming agents that increase total input mass, such as those presented here, may not be thought to improve the performance of these assays. However, as our results with DNA1 suggest, it may be possible to develop agents that specifically enrich informative subsets of fragments of cfDNA for these assays.

In summary, this study has extended antibody-based priming agents for liquid biopsies from a single proof-of-concept mAb to a wide range of agents, uncovering key features that govern the priming effect and revealing new insights into the biology of cfDNA. The finding that non-immunoglobulin binding domains can also have a priming effect supports the generalizability of this approach beyond antibodies. Future work leveraging these principles could lead to more efficient screening and rational design of priming agents to maximize their priming effect and advance investigations into the biology of cfDNA.

## Acknowledgements

This work was supported by a Cancer Center Support (core) grant P30-CA14051 from the National Cancer Institute, the Gerstner Family Foundation (VAA), the Koch Institute Frontier Research Program via the Casey and Family Foundation Cancer Research (JCL, SNB), the Bridge Project, a partnership between the Koch Institute for Integrative Cancer Research at MIT and the Dana-Farber/Harvard Cancer Center (JCL, VAA), and the Koch Institute’s Marble Center for Cancer Nanomedicine (SNB). ST acknowledges support from NIH (K08EB036081), the Prostate Cancer Foundation (Young Investigator Award 21YOUN20), and Massachusetts General Hospital Department of Radiation Oncology. CMA acknowledges support from a fellowship from “La Caixa” Foundation (ID 100010434). The fellowship code is LCF/BQ/AA19/11720039. CMA also acknowledges support from The Ludwig Center Fellowship at MIT’s Koch Institute. DMK was supported by award Number T32GM007753 and T32GM144273 from the National Institute of General Medical Sciences and award number F30CA298604 from the National Cancer Institute. The content is solely the responsibility of the authors and does not necessarily represent the official views of the National Institute of General Medical Sciences or the National Institutes of Health. S.N.B. is a Howard Hughes Medical Institute Investigator.

## Materials and Methods

### Antibody Design and Expression

Antibody priming agents were designed using published VH and VL sequences for antibodies reported to bind dsDNA, histones, or nucleosomes.^11–15^ Sequences were prioritized based on completeness, extent of characterization of the mAb in associated publication, and reported binding strength to the target antigen. VH sequences were fused to mIgG2a Fc sequence with the L234A/L235A/P329G silencing mutations.^10^ Eighteen candidate antibodies were cloned and expressed using the TurboCHO system (Genscript) with one-step protein A purification into PBS (pH 7.2) followed by downstream quality control via SDS-PAGE (reduced and non-reduced), SEC-HPLC, and endotoxin testing (Genscript TurboCHO or High Throughput TurboCHO platforms). Sso7d-Fc fusions were designed and produced using the same system and all carried the same L234A/L235A/P329G mutations. Mono-sso7d was produced in single arm format by heterodimerization with a truncated Fc domain using “hole” (T366S/L368A/Y407V) and “knob” (T366W) mutations in the sso7d-Fc and truncated Fc sequences, respectively.^39^ Antibody preparations were maintained at −80C until use. Antibody concentration was determined by nanodrop A280 (Denovix DS-11) upon thawing.

### Electrophoretic Mobility Shift Assay (EMSA)

Widom601 dsDNA complexed with recombinant human histones was purchased from Epicypher (cat. 16-0009). Free Widom601 dsDNA was amplified from a purchased template (cat. 18-0006, Epicypher) using primers 5’-CTGGAGAATCCCGGTGC and 5’-ACAGGATGTATATATCTGACACGTGC. Free and histone bound dsDNA were mixed to a final dsDNA concentration of 2 ng/μL and 1 ng/μL, respectively, with antibodies at a final concentration of 100nM in a filtered reaction buffer (10mM Tris, 50mM NaCl, 1mM DTT, 0.1 mg/ml BSA, 6% glycerol, 1mM EDTA, 0.5X TBE) to 10 μL total. The mixture was incubated for 30 minutes at RT followed by the addition of 1 μL of 5X Novex Hi-Density TBE Sample Buffer (cat. LC6678, ThermoFisher). 10 μL of the sample was added to a 6% polyacrylamide DNA retardation gel (cat. EC63652BOX, ThermoFisher) and the gel was run at 4°C for 2 hours at 100V in 0.5X TBE running buffer. The gel was stained with 1:1000 SYBR Safe (cat. S33112, ThermoFisher) in 0.5X TBE for 30 minutes and imaged with an iBright imager (ThermoFisher).

### Biolayer Interferometry

Streptavidin biosensor tips (cat. 18-5019, Sartorius) were used to immobilize mononucleosomes (16-0006, Epicypher), free dsDNA (18-0005, Epicypher) with the Widom601 sequence and a biotin tag, or biotinylated supercoiled or linearized pUC18 plasmid DNA, diluted to a final concentration of 8nM in kinetics buffer (5mg/ml BSA, 0.05% Tween-20 in PBS). Interaction with mAbs, diluted to various concentrations in filtered kinetics buffer, was tested at 37°C using Sartorius Octet R8 according to the following assay details: Baseline 150s, Loading 500s, Baseline2 100s, Association 400s, Dissociation 400s. Kinetics recordings were standardized to the start of the association phase and kinetic parameters and K_d_ values were estimated using best fit nonlinear regression in GraphPAD Prism.

For biotinylated antibodies, BLI was performed by immobilizing the antibodies diluted in kinetics buffer (5mg/ml BSA, 0.05% Tween-20 in PBS) on Streptavidin biosensor tips (cat. 18-5019, Sartorius). Binding to histone proteins diluted in kinetics buffer was measured on a Sartorius Octet R8 at 37C according to the following assay details: Baseline 150s, Loading 500s, Baseline2 100s, Association 400s, Dissociation 400s.

### Biotinylation of antibodies and plasmids

Antibodies at a concentration of 0.8 mg/mL were incubated at a 1:1 molar ratio with EZ-Link NHS-PEG4-Biotin (cat. A39259, ThermoFisher) in PBS at RT for 30 minutes and buffer exchanged into PBS using Zeba spin desalting columns (7K MWCO, cat. 89882, ThermoFisher). pUC18 plasmid (cat. SD0051, ThermoFisher) was biotinylated using the Label IT siRNA Tracker kit (cat. MIR7217, MirusBio). 20 μL of pUC18 plasmid DNA at 0.5 μg/μL was combined with 10 μL of 10X labeling buffer A, 2.5 μL of Label IT reagent and 67.5 μL of molecular biology-grade water. The reaction was incubated at 37°C for 1 hour. Plasmid DNA was purified using ethanol precipitation. 10 μL of 3M sodium acetate was added to the reaction followed by 275 μL of 100% ethanol at −20°C. The reaction was incubated at −80°C for 1 hour and centrifuged at 20,000g at 4°C for 30 min. The supernatant was removed, the pellet washed with 250 μL of chilled 70% ethanol, and centrifuged at 20,000g at 4°C for 5 min. The supernatant was removed and the pellet was allowed to dry before being resuspended in water. Biotinylated pUC18 plasmid was linearized using BamHI (cat R0136S, NEB) per manufacturer instructions with 3h incubation at 37°C and purified using the QIAquick PCR purification kit (cat. 28104, Qiagen). Linearization was confirmed with agarose gel electrophoresis (E-gel EX 1%, cat. G401001, ThermoFisher).

### F(ab’)2 preparation

Antibody without LALAPG Fc mutations was used for F(ab’)2 preparation as mutations conferred protease resistance. IdeZ protease (cat. V8341, Promega) was added to 1mL of antibody in PBS at a ratio of 1μL IdeZ: 50 μg antibody and incubated at 37°C for 2 hours. The reaction was cooled to room temperature and IdeZ was removed by adding MagneHis Ni-Particles (cat. V8560, Promega) at a ratio of 30 μL Ni particles: 1 mL IdeZ protein reaction. The Ni particles were mixed thoroughly and incubated for 2 minutes at room temperature, followed by magnetic separation and removal of the supernatant containing Fc and F(ab’)2 fragments. The resulting fragmented sample volume was measured and used to calculate the volume of Protein A Beads (cat. G8781, Promega) that would be used to isolate the F(ab’)2 at a ratio of 1 μL beads: 2 μL sample. Protein A beads (cat. G8781, Promega) were prepared by washing with 0.5X the volume with PBS, followed by two additional 1mL PBS washes. The beads were then resuspended in the sample and incubated at room temperature for 30 minutes with shaking at 1000 rpm. The supernatant containing the purified F(ab’)2 was collected using magnetic separation. Purification was verified on a protein gel and the concentration of resulting F(ab’)2 was measured using A280 on a Nanodrop spectrophotometer (Denovix DS-11) with E1% 14.0.

### Nuclease protection assay

Free dsDNA or MN (cat. 16-0009, 18-0006 respectively, Epicypher) was prepared at final DNA concentration of 4ng/uL and incubated with mAbs at 400nM at 37°C in 37 μL DNase buffer (B0303S, New England Biolabs) for 5 min. Following incubation, 1uL DNase I (M0303S, New England Biolabs) at 7.6U/mL was added to a final concentration of 0.2U/mL, consistent with the reported range of nuclease activity in plasma.^21–23^ Immediately after addition of DNase and at subsequent time intervals, 11 μL of the reaction mixture was mixed with 1.5 μL of 25 μM EDTA to stop the reaction. 1 μL of 20mg/mL proteinase K (Qiagen cat. 19134) and 1.5 μL of 5% SDS were added to the reaction. The reaction was incubated at 55°C for 30 minutes and then cooled to room temperature. The reaction was cleaned using a 2X ratio of AMPure XP (Beckman Coulter) and eluted in 15 μL of lowTE buffer. DNA concentration was determined using the Quant-iT HS assay (cat. Q33232, ThermoFisher) according to manufacturer instructions.

### Animal Models

All animal studies were approved by the Massachusetts Institute of Technology Committee on Animal Care (MIT Protocols 042002323, 2301000462). Animals were maintained in the Koch Institute animal facility with a 12h-light/12h-dark cycle at 18-23°C and 50% humidity. Female BALB/c mice (6-10 weeks, Taconic Biosciences) were used for all healthy mice experiments. For tumor-bearing mice, the murine colon cancer cell-line Luc-MC26 (expressing firefly luciferase, gift from the Kenneth K. Tanabe Laboratory) was used. Cells were cultured in ATCC-formulated Dulbecco’s Modified Eagle’s Medium DMEM (cat:10-013-CM, Corning) supplemented with 10% fetal bovine serum (cat 100-500, GemCell) and 1% penicillin/streptomycin (cat 30-002-C1, Corning), in a humidified atmosphere of 95% air and 5% CO_2_ at 37°C. To establish lung tumors, 1×10^5^ Luc-MC26 cells were injected intravenously i.v. into female BALB/c mice (6 weeks, Taconic Biosciences). Tumor burden was monitored by luminescence using the In Vivo Imaging System (IVIS, PerkinElmer) every 3-5 days after tumor inoculation.

### In vivo cfDNA pharmacokinetics

Widom601 dsDNA, either free (cat. 16-0006, Epicypher) or histone bound in mononucleosomes (18-0005, Epicypher), was mixed to a final concentration of 0.2 ng/μL with each mAb binder to a concentration of 0.4 mg/mL in DPBS. Each mouse was anesthetized with inhaled isoflurane and injected i.v. with 200 μL of mixture. At 1 minute and 60 minutes post injection, 75 μL of blood was collected via a retro-orbital blood draw, both under anesthesia. Mice were allowed to recover from anesthesia between blood draws. Widom601 quantification was performed as described below.

### Testing priming agents in tumor-bearing mice

Priming agents were tested in tumor-bearing mice approximately 14 days after tumor inoculation. Priming agents were thawed, the concentration was measured via A280, and the agent was diluted to 0.4 mg/mL with sterile PBS. 75 μL of blood was sampled retro-orbitally from each mouse prior to treatment. Prior to administration, the priming agent preparation was sterile-filtered (cat. 9916-1302, Whatman). 4.0 mg/kg of antibody priming agents or control PBS were administered into awake mice i.v. Two hours following administration, 75 μL of blood was collected retro-orbitally from the contralateral eye, consistent with our previous work on priming agents.^9^ cfDNA concentration measurement and ctDNA detection was performed as described below.

### Sample processing

75 μL retro-orbital blood draws were collected with nonheparinized capillary tubes from mice under isoflurane anesthesia, alternating between eyes for serial draws. Blood was immediately displaced from the capillary tube into 75 μL of 10-mM EDTA (cat. AM9260G, Thermo Fisher Scientific) in PBS. Blood with EDTA was kept on ice and centrifuged within 60 min of collection at 8000g for 5 min at 4°C. The plasma fraction was collected and stored at –80°C until further processing

### cfDNA extraction and quantification

Frozen plasma was thawed and centrifuged at 15000g for 10 minutes for pellet separation. Plasma was combined with PBS to a total volume of 2100 μL and cfDNA was extracted using the QIAsymphony DSP circulating DNA kit (cat. 937555, Qiagen) according to the manufacturer’s instructions, with an elution volume set to 60 μL. Endogenous extracted cfDNA was quantified using Taqman qPCR targeting a locus on chromosome 8 in the mouse genome (forward primer: GGGACTCCTGCAGATCGTTA, reverse primer: ATCTGGCCCTATCTTCCATCCT, Taqman probe: /56-FAM/CCTGTGGTG/ZEN/CTGAACCTATCAACAGCA/3IABkFQ/). Exogenously delivered Widom 601 DNA was similarly quantified using Taqman PCR targeting the Widom 601 sequence (forward primer: 5’CGCTCAATTGGTCGTAGACA, reverse primer: 5’TATCTGACACGTGCCTGGAG and Taqman probe: /56-FAM/TCTAGCACC/ZEN/GCTTAAACGCACGTA/3IABkFQ/). % W601 remaining was calculated as the % W601 remaining at 60 min relative to 1 min. Extracted DNA was kept at 4°C until further processing.

### Immunoprecipitation of cfDNA

Antibodies were covalently coupled to Dynabeads M-270 Epoxy beads (cat. 14311D, Thermo) using the manufacturer’s protocol at a ratio of 6 μg antibody to 1 mg beads. 20 μL of resulting mAb-coupled beads were pre-washed with PBS + 0.1% BSA for 5 minutes and resuspended in 20 μL PBS, then mixed with 100 μL of BALB/c Mouse plasma (cat. IGMSBCPLAK2E50ML, Innovative Research) in PBS 5mM EDTA (cat. 15575020, IDT) to 130 μL. For MN pulldown, experiments, 5 ng of Widom601 free DNA (18-0006, Epicypher), 5 ng of Widom601 MN (16-0009, Epicypher) or 10 ng of DNase-treated Widom601 MN were added to the plasma sample. To achieve partial digestion of free DNA in Widom 601 MN, DNA was mixed with DNase I (EN0521, Thermo) at 0.25 U DNase per 1 μg MN, incubated at 37°C for 10 minutes, then quenched with EDTA to 15mM. Following plasma sample preparation, reactions were incubated at RT for 15 minutes with shaking at 800 rpm. Beads were separated using a magnet, and the supernatant was collected and stored at RT until further processing. Beads were washed 3X with 200 μL of PBS (off magnet), and eluted in 110 μL of PBS with 5mM EDTA. Bead and supernatant fractions were processed using standard DNA extraction and cfDNA quantification. For experiment using Protein A/G immunoprecipitation, 1 μg antibody was added directly to BALB/c Mouse plasma, mixed with PBS 5mM EDTA to 110 μL. 30 μL of Protein A/G beads (cat. 88803, ThermoFisher) were pre-washed with 90 μL and 200 μL of 1X TBST (cat J77500.K2, ThermoFisher). Following a 15 minute RT incubation of the sample, dried beads were resuspended in the plasma mixture and incubated on a rotating mixer for 1 hour. Supernatant and bead fractions were separated and processed as described above, with the substitution of a 3X bead wash with 200 μL of 1X TBST prior to bead elution.

### Pharmacokinetic study of Fc-modified antibodies

To label antibodies, the amine-reactive fluorophore AQuora® 750 (cat. AQ-11960LF, Quanta Biodesign) was incubated with 450μL aliquots of 2 mg/mL mAb (or 1.6 mg/mL F(ab’)2) in PBS, at 15 dye: 1 protein molar ratio, for 2h at room temperature, protected from light. Excess dye was removed by washing conjugate with sterile PBS 9 times using 30kDa Amicon filters (cat. UFC503024, EMD Millipore) spun at 15,000g for 3.5min at 4C. After concentrating the volume of the last wash to approximately 200 μL, a Nanodrop spectrophotometer (Denovix DS-11) was used to measure the protein yield (absorbance at 280nm) and the fluorophore concentration and degree of labelling (absorbance at 755nm, A280 correlation 0.04, background correction at 840nm). Degree-of-labeling ranged from 3.02 to 7.07. Labeled proteins were kept at 4C and protected from light until further use. Prior to animal studies, labeled antibodies were sterile filtered and requantified. For antibody pharmacokinetic studies, AQuora® 750-labeled antibodies were injected i.v. at 4.0 mg/kg (200μL in sterile PBS) into anesthetized mice and 75μL of blood was collected from alternating eyes at 1min, 2h, 6h and 24h. Abundance of antibodies in plasma samples was measured using a Tecan fluorometer at 750/785nm and a standard prepared with unconjugated dye, then calculated as % remaining relative to the 1 min plasma sample.

### Library Construction, Hybrid Capture, Sequencing

cfDNA libraries were constructed from extracted cfDNA using the KAPA Library Prep kit (cat. KK8504, Kapa/Roche), with custom dual index UMI adapters (IDT) and indexing primers (cat. 10009816, IDT). For whole genome sequencing (WGS), libraries were sequenced to a target depth of 2x (range 1-4x) on an Illumina NovaSeq S4 or NovaSeqX (151 bp paired-end reads). For the ctDNA diagnostic test, two rounds of hybrid capture using a tumor specific panel were done on the libraries using the xGen hybridization and wash kit (cat. 1080584, IDT), xGen Universal blockers (cat. 1075476, IDT), and a previously-described custom tumor specific panel targeting 1822 SNVs in the MC26 murine colon cancer cell line.^9^ Libraries were pooled up to maximum 18-plex, with a library mass equivalent to 25x DNA mass into library construction for each sample, and the previously used panel consisting of 120-bp long probes (IDT) targeting tumor-specific SNVs was applied.^9^ After the first round of HC, libraries were amplified by 16 cycles of PCR and then carried through a second HC. After the second round of HC, libraries were amplified through 8 to 16 cycles of PCR, quantified, and then pooled for sequencing. Pooled HC libraries were sequenced to a targeted raw depth of 50,000X per site per 20 ng of cfDNA input into library construction on Illumina NovaSeq S4 or NovaSeqX (151 bp paired-end reads). Sequencing data were processed using a duplex consensus calling pipeline as previously described,^9^ yielding measurements of the total number of mutant duplexes detected, the unique number of loci detected, and the tumor fractions.

### Analysis of coverage and end-motifs

Demultiplexed WGS FASTQ files for cfDNA samples were aligned to the mouse genome (mm9). Coordinates of mouse CpG islands were downloaded from UCSC genome browser (table cpgIslandExt, assembly mm9). For promoter regions, we took 1kb upstream and 100bp downstream of transcription start sites (table knownGene, assembly mm9). We downloaded DNase HS peaks for all leukocyte and myeloid datasets in mouse ENCODE (ENCFF063EHX, ENCFF125IXR, ENCFF171XTE, ENCFF185ZCW, ENCFF215CPX, ENCFF359GEV, ENCFF434GOV, ENCFF550NKM, ENCFF566TDU, ENCFF689PKR, ENCFF702OKE, ENCFF754XGR, ENCFF761OVL) and converted to mm9 coordinates using LiftOver (https://genome.ucsc.edu/cgi-bin/hgLiftOver). ENCSR000CMQ and ENCSR000CNP were excluded due to very low read depth. We also generated 100 20-kb random regions in the genome as control regions. Aligned bam files were filtered to separately create bam files with template length <=120 bp and bam files with template length > 120bp. In each filtered bam file, raw coverage depth was calculated using mosdepth^46^ for each set of genomic features (CpG islands, promoters, DHS) and normalized to genome-wide depth of coverage to obtain relative coverage in each feature. We then calculated the ratio of relative coverage by fragments <=120bp and >120bp for each feature as a measure of fragment length dependent coverage bias.

To identify 4-mer endmotifs from each fragment, we first needed to remove UMI sequences from each read. Since our adapter pool consisted of a 1:1 mixture of 3bp and 4bp UMIs and a T overhang (“UMI+T”), we could not use a fixed length to eliminate UMI’s: trimming 5bp would remove an additional base from 50% of reads whereas trimming 4bp would keep the T overhang from the adapter in 50% of reads. To confidently trim UMI+T, we first realigned each FASTQ without including UMI or T overhang in the read structure. We then filtered to only include reads that had 5 soft-clipped bases at the 5’ end followed by at least 20 perfectly aligned bases and overall read MAPQ > 20. This filter would keep only those reads with 4bp UMIs followed by a T overhang, allowing us to then trim 5bp to find the starting sequence of the template. To confirm this, we checked the last soft-clipped base of our filtered reads, which should always be a T (representing the T overhang on the adapter), and found that 99.4% of reads had a T in this position. The remaining 0.6% likely represents errors introduced during PCR and sequencing. Having successfully identified the true start position of the read, we then extracted the first 4-bp from the 5’-end of the template. 3’-end sequences were not used due to 3’ exonuclease processing during end repair. Frequencies of 4-mer end-motifs in cfDNA in each sample were normalized based on observed frequency across the mouse genome, tabulated using Jellyfish^47^. End-motif frequencies were compared across priming agents using two-tailed t-tests, p-values were adjusted using the Benjamini-Hochberg procedure, and motifs with adjusted p < 0.01 and log2(fold-change) > 0.5 or < −0.5 were considered significant. For analysis of F-profile contribution, we used non-negative least squares from *scipy.optimize* to fit the observed normalized frequencies in each sample to the F-profiles reported by Zhou et al.^26^

Griffin^29^ was used for analysis of nucleosome patterns in cfDNA. For transcription factor binding sites, we used meta clusters (downloaded from https://gtrd.biouml.org/downloads/19.10/chip-seq/Mus%20musculus_meta_clusters.interval.gz), which contains meta peaks in one or more ChIP seq experiments. We selected the top 5000 binding sites by peak signal for each transcription factor and changed coordinates from mm10 or mm9 using LiftOver (UCSC genome browser). Mappability and blacklist files were downloaded from https://hgdownload.cse.ucsc.edu/goldenPath/mm9/encodeDCC/wgEncodeMapability/ and http://mitra.stanford.edu/kundaje/akundaje/release/blacklists/mm9-mouse/mm9-blacklist.bed.gz respectively. Griffin was run with GC correction and default parameters, except with modification of fragment lengths analyzed for specific analyses, as noted.

### Statistical analysis

A suite of scripts (Miredas) was used for calling SNVs and creating metrics files. All other analyses were performed using GraphPad Prism v9, custom Python scripts and R. Unpaired two-sided t-tests were used for comparisons unless noted otherwise. For each tumor-bearing animal experiment, mice were allocated such that groups would have comparable tumor burden. Investigators were not blinded to the groups and treatments during the experiments.

