## Supplementary Figures for "Molecular determinants of antibody-mediated priming to enhance detection of ctDNA"

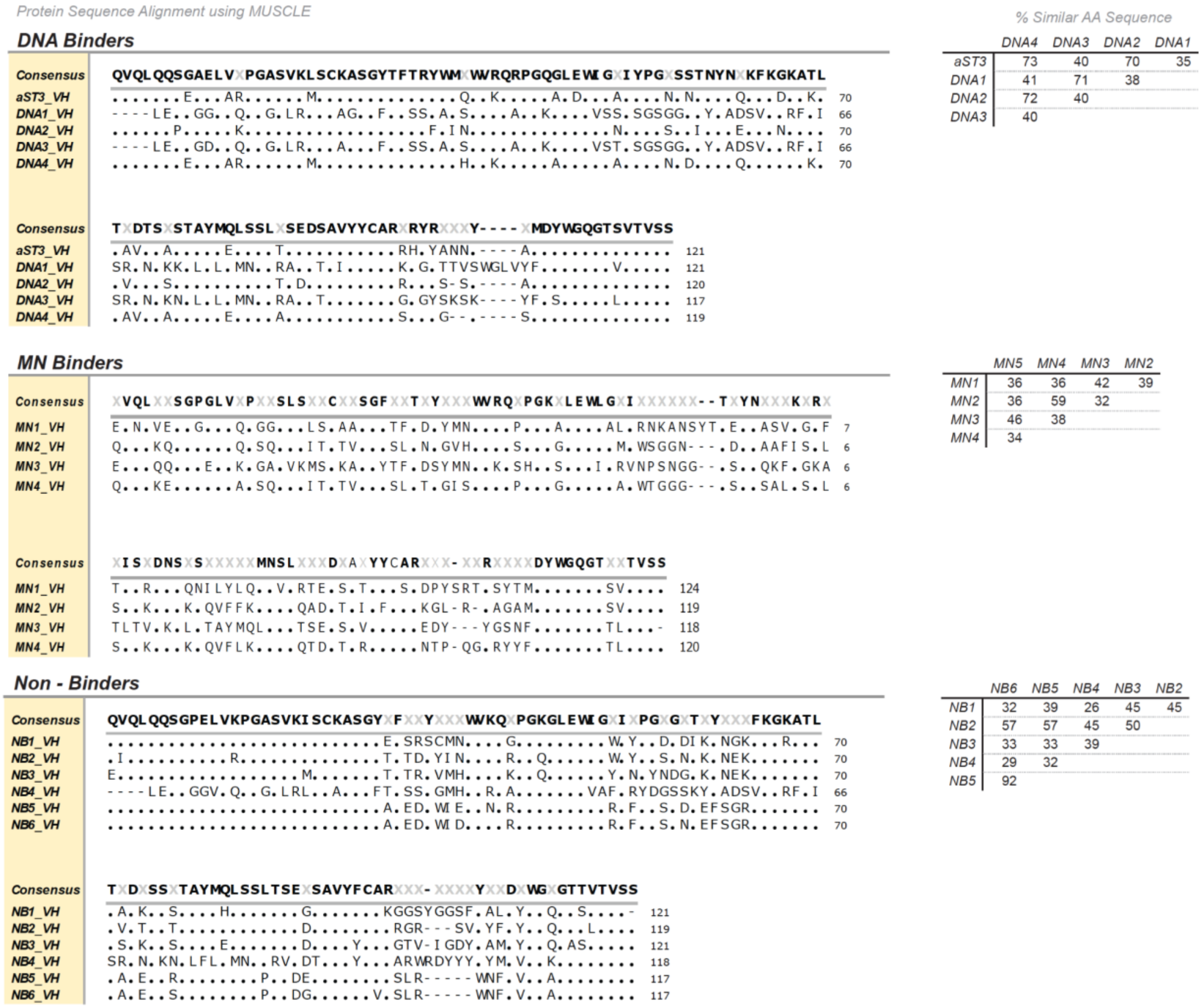


**Figure S1:** Protein sequences of heavy chain variable domains for aST3 and the 14 other mAbs aligned using MUSCLE. Sequence alignment is separated by binding classification, and percent sequence similarity between the mAbs is calculated within the binder groups.

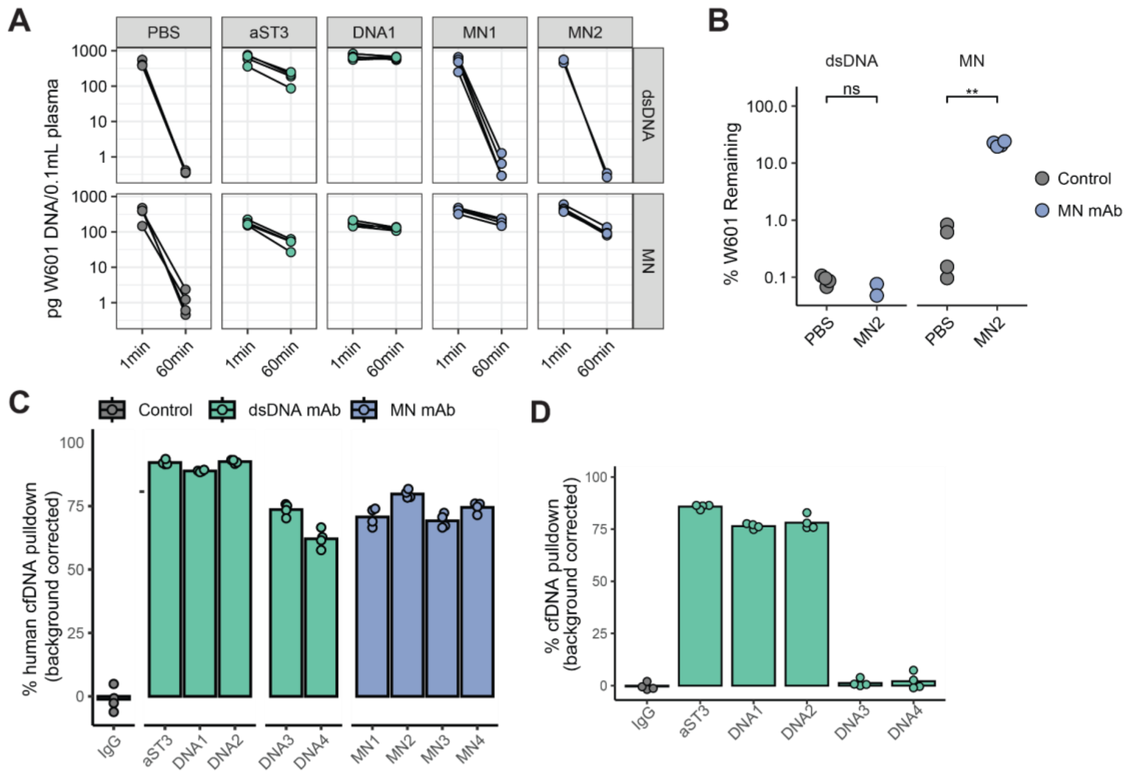


**Figure S2: Clearance and immunoprecipitation of exogenous and endogenous cfDNA using mAbs.** A) Mass of either free dsDNA or MN bound W601, normalized to 0.1mL of plasma, detected in blood draws 1 minute and 60 minutes post-injection. B) Percent of dsDNA and MN W601 remaining at 60 minutes, relative to 1 minute recording, for MN2 and PBS. C) Percent of cfDNA mass pulled down from human plasma using mAb-coupled magnetic beads. D) Percent of cfDNA mass, relative to the total collected sample mass, pulled down from mouse plasma with addition of mAb priming agents followed by addition of protein A magnetic beads.

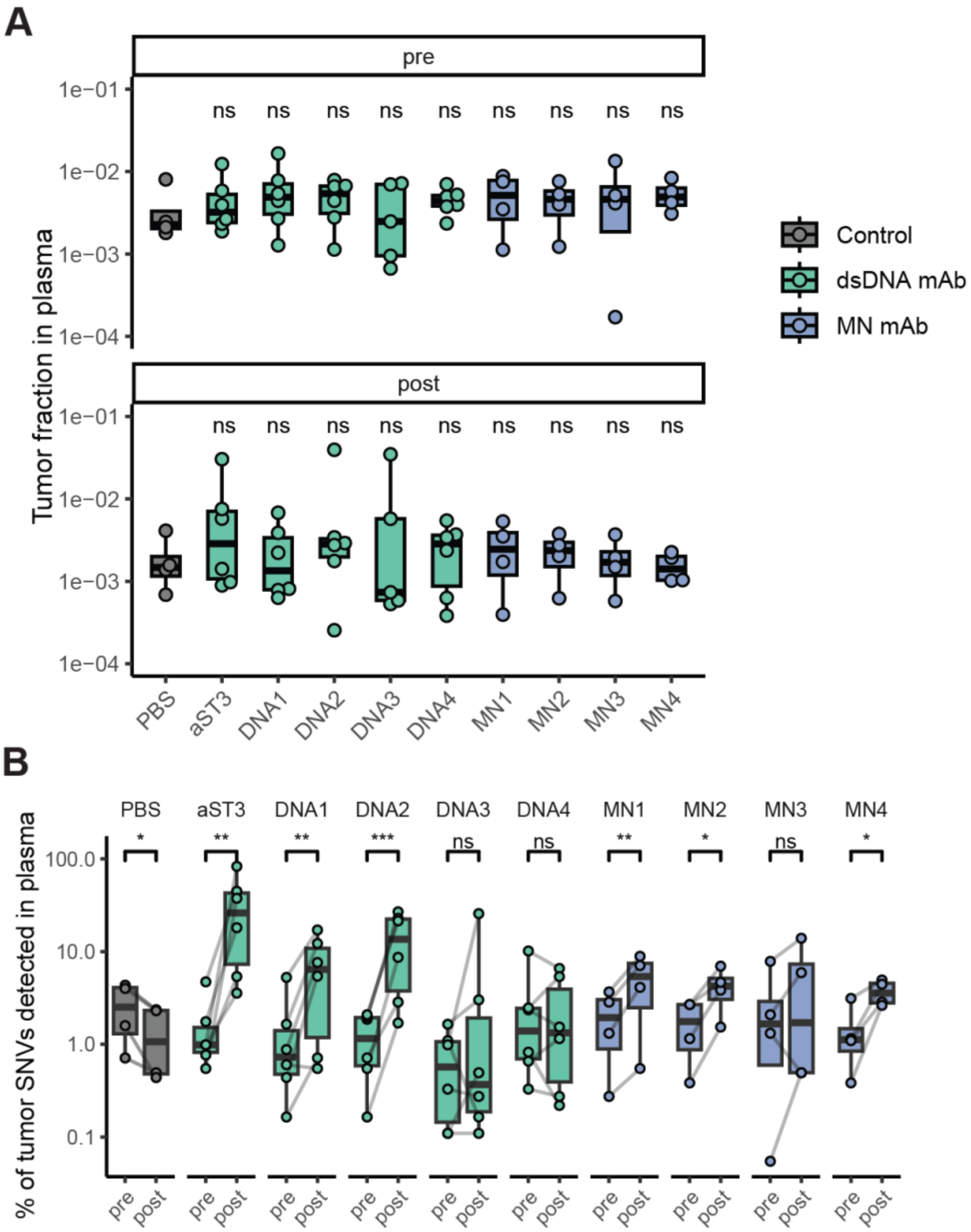


**Figure S3: Tumor fraction and SNV detection pre and post mAb priming.** A) Tumor fraction before ("pre") and 2 hours after ("post") administration of priming agents, where tumor fraction represents the ratio of ctDNA molecules to total cfDNA molecules. ns - not significant. B) Percentage of 1,822-SNV tumor mutation panel that was detected in plasma before and after administration of priming agents.

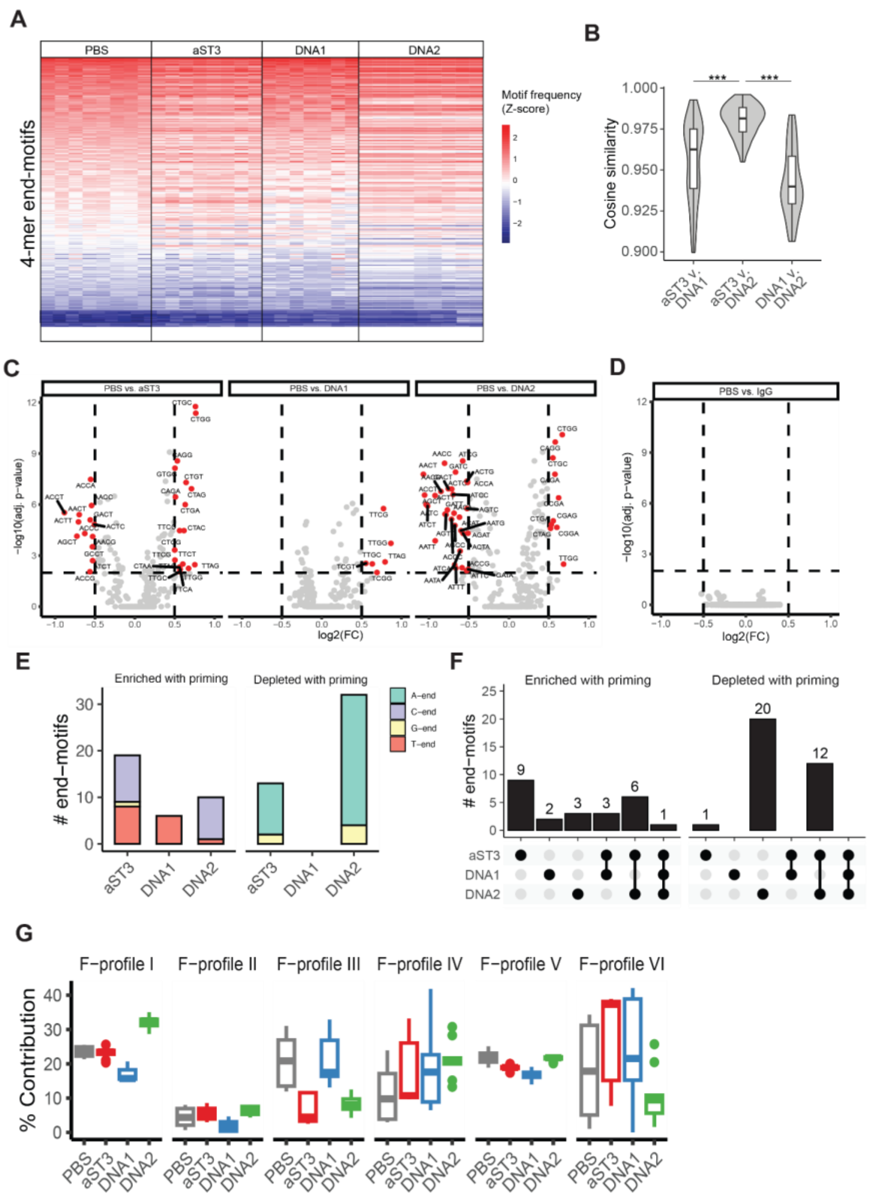


**Figure S4: End motif analysis after priming.** A) 4-mer end-motif profiles after administration of priming agents aST3, DNA1, and DNA2, versus PBS control. B) Cosine similarity of 4-mer end-motif profiles after administration of different priming agents. C) Comparison of 4-mer end-motif frequency after administration of DNA-binding mAb priming agent versus PBS control and D) after administration of IgG control mAb versus PBS control. End-motifs that have higher frequency in priming agent samples have log2(FC) > 0 and those that have higher frequency in PBS samples have log2(FC) < 0. End-motifs significant at adjusted p-value < 0.01 with log2(fold change) in frequency greater than 0.5 or less than -0.5 are highlighted. E) Number of 4-mer end-motifs that are significantly enriched or depleted after priming agent administration compared to PBS samples. Colors indicate the first base in the motif, at the fragment end. F) Number of significantly enriched and depleted 4-mer end-motifs that are unique to each priming agent or shared across priming agents. G) Contribution of F-profiles I-VI to the end-motif profiles of plasma cfDNA after PBS, aST3, DNA1, and DNA2.

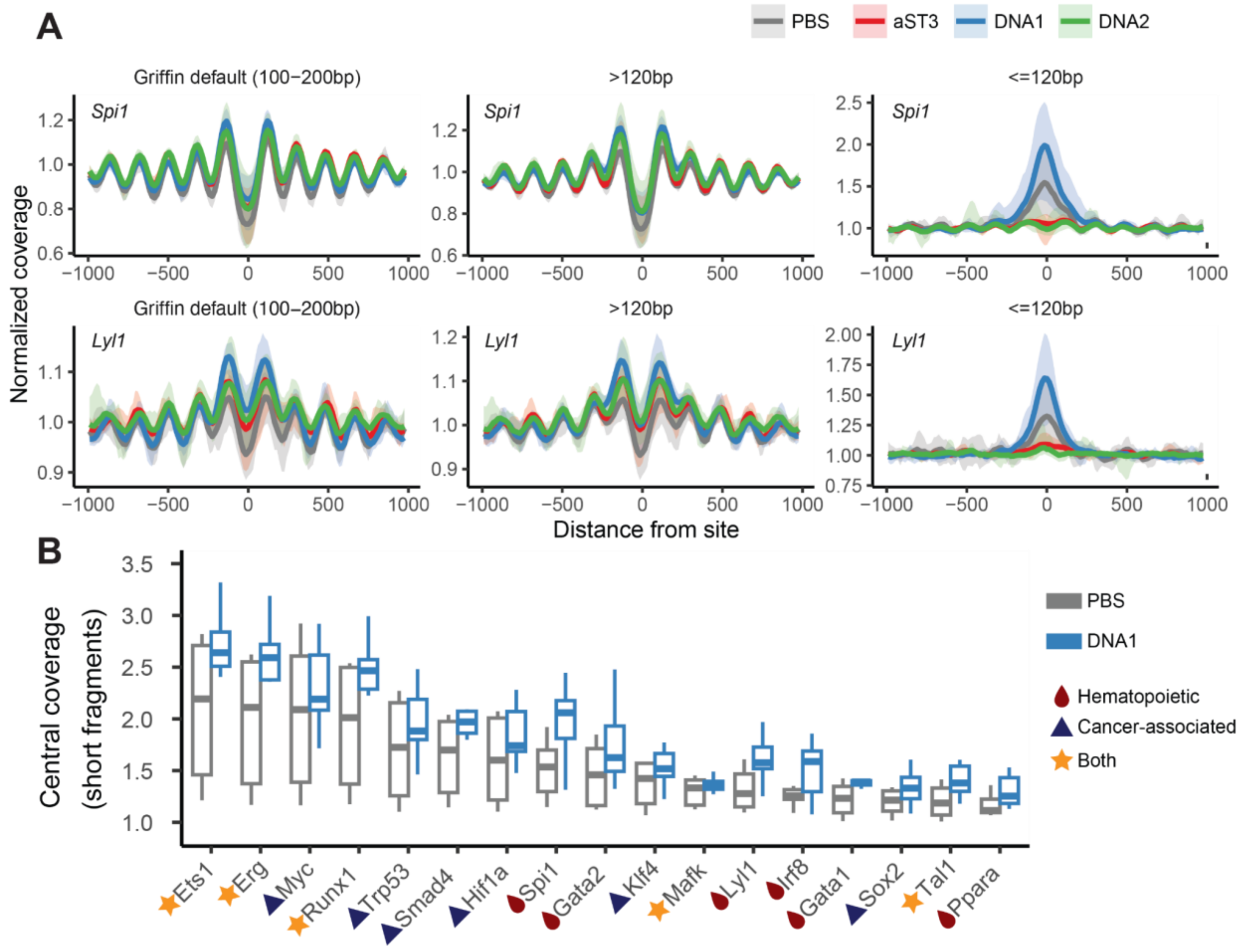


**Figure S5: cfDNA coverage patterns after administration of priming agents.** A) Coverage patterns after administration of aST3, DNA1, DNA2, or PBS control, around Spi1 and Lyl1 binding sites using Griffin's default fragment filter (100-200bp), long fragments (>120bp, range 121-500bp), and short fragments (<= 120bp, range 15-120bp). Bands indicate the range across samples for each agent. B) Central coverage across a range of hematopoietic and cancer-associated transcription factor binding sites by short fragments (<=120 bp) in cfDNA after administration of PBS control or DNA1.

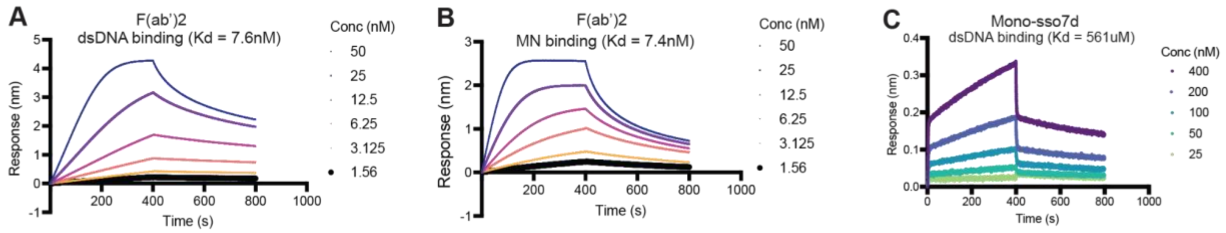


**Figure S6: Biolayer interferometry traces of engineered priming agents at varying concentrations.** A) Binding of aST3 F(ab')2 to immobilized dsDNA. B) Binding of aST3 F(ab')2 to immobilized mononucleosomes (MN). C) Binding of mono-sso7d to immobilized free dsDNA.
